# Cell types in the mouse amygdala and their transcriptional response to fear conditioning

**DOI:** 10.1101/2022.10.25.513733

**Authors:** Hannah Hochgerner, Muhammad Tibi, Shai Netser, Osnat Ophir, Nuphar Reinhardt, Shelly Singh, Zhige Lin, Shlomo Wagner, Amit Zeisel

## Abstract

The amygdala is one of the most widely studied regions in behavioral neuroscience. A plethora of classical, and new paradigms have dissected its precise involvement in emotional and social sensing, learning, and memory. Several important insights resulted from the use of genetic markers – yet, in the age of single cell transcriptomics, the amygdala remains molecularly underdescribed. Here, we present a molecular cell type taxonomy of the full mouse amygdala in fear learning and consolidation. We performed single-cell RNA-seq on naïve and fear conditioned mice, inferred the 130 neuronal cell types distributions in silico using orthogonal spatial transcriptomic datasets, and describe the cell types’ transcriptional responses to learning and memory consolidation. Only a fraction of cells, within a subset of all neuronal types, were transcriptionally responsive to fear learning, memory and retrieval. These activated engram cells upregulated activity-response genes, and processes of synaptic signaling, plasticity, development and neurite outgrowth. Our transcriptome-wide data confirm known actors, and describe several new candidate genes. The atlas may help pinpoint the amygdala’s circuits in performing emotional sensing and integration, and provide new insights to the global cellular processes involved.

## Introduction

The amygdala is part of the medial temporal lobe of the brain. It consists of multiple anatomical regions, developmentally attributed to the striatal, cortical (superficial), and subcortical (deep) structures^1^. These subregions are defined by their distinct cytoarchitecture, connectivity, and functionality. Among the commonly studied amygdala functions is fear learning and its circuitry, which involves basolateral, intercalated, and central amygdala clusters. Learning paradigms reflecting other, non-aversive valences (e.g., appetitive behaviors) have since revealed more nuanced amygdala roles in emotional sensing. These alternate functions were often achieved through neuronal populations in the same regional circuitries, distinguished only by different molecular identities^2–4^.

Classified as part of the olfactory system, other amygdala regions are widely studied in the social and emotional contexts, where they coordinate adaptive behavioral responses. For example, the medial amygdala is involved in social, parenting, and aggression behaviors, and in memory tasks such as social recognition (reviewed in Petrulis, 2020). Several medial amygdala functions are linked to the gonadal steroid sensitivity of its neurons, some of which have been described systematically^6^. The cortical amygdala receives direct input from the olfactory bulb, where spatially distinct neurons are activated by odors of positive or negative valence^7,8^. Finally, neurons of the basomedial amygdala participate in aspects of fear learning as well as defensive and social behaviors^9,10^. This region has very limited molecular description^11^.

Functional studies have used molecular markers, such as *Fezf2*, *Grp*, *Rspo2* and *Ppp1r1b* in LA/BLA, *Prkcd* in CeA, *Foxp2* in IA, and *Trh* in CoA,^3,4,12–14^. But cells defined by a single selected marker gene do not necessarily correspond to a unique neuron type. Four studies described murine amygdala cell types in detail, using scRNA-seq of cells isolated from the medial amygdala^6,15^, basolateral amygdala^16^ and full amygdala part of a cell type atlas of the mouse nervous system^17^. The studies described molecular cell types in details exceeding those of the markers previously known, and suggested a high diversity of underexplored cell types in this brain region. In addition, single-cell transcriptomics has been used for detecting transcriptional signatures of neuronal activation^18,19^, social behaviors^20,21^, and learning and memory^22,23^.

Aiming to expand the “parts list” available to researchers working in this dynamic field, we used unbiased, high-quality scRNA-seq on whole cells gently dissociated from full amygdala in 23 adult, naïve and fear conditioned mice. We applied parallel approaches to infer spatial distributions of the resulting taxonomy to uncover molecular principles governing this unique niche of developmentally and functionally distinct regions. Finally, we describe which neuronal types participated in fear learning, and the orchestrated transcriptional response across consolidation and retrieval.

## Results

### Whole-cell scRNA-seq on amygdala of adult mice

We refined our dissociation protocol to optimize cell viability for conducting whole-cell scRNA-seq on amygdala of adult mice, a region we previously found to be particularly sensitive to dissociation stress^17^. As for other cell dissociations from brain tissue, it was important to minimize handling time from the sacrificing of the mice to loading the cells for scRNA-seq, as well as any stress experienced by the cells during this time (temperature, enzyme, mechanical; see Methods). Significant improvements in viability came from using a single buffer (artificial cerebrospinal fluid, aCSF), which was formulated particularly for the more sodium-sensitive neurons of adult and aging animals^24^ (see Methods).

We verified improved amygdala neuronal viability in suspension, which resulted in high scRNA-seq data quality (e.g., >3000 UMI, >2500 genes per neuron). We performed scRNA-seq on full amygdala of 23 adult mice, 16 of which had undergone tone-cued fear conditioning (CFC, see Methods, Discussion, Suppl. Table 1) (Fig. 1a). Preliminary analysis revealed 25,330 non-neuronal cells that organized into 14 clusters (Fig.1b, Extended Data Fig. 1), and great diversity among 30,184 high-quality neurons. Neurons were robustly segregated based on their neurotransmitter identity to GABA (*Gad1, Gad2, Slc32a1*) or GLUT, and the glutamatergic ones, into dominant vesicular glutamate transporter (VGLUT) 1 (*Slc17a7*) or 2 (*Slc17a6*).

**Figure 1.**
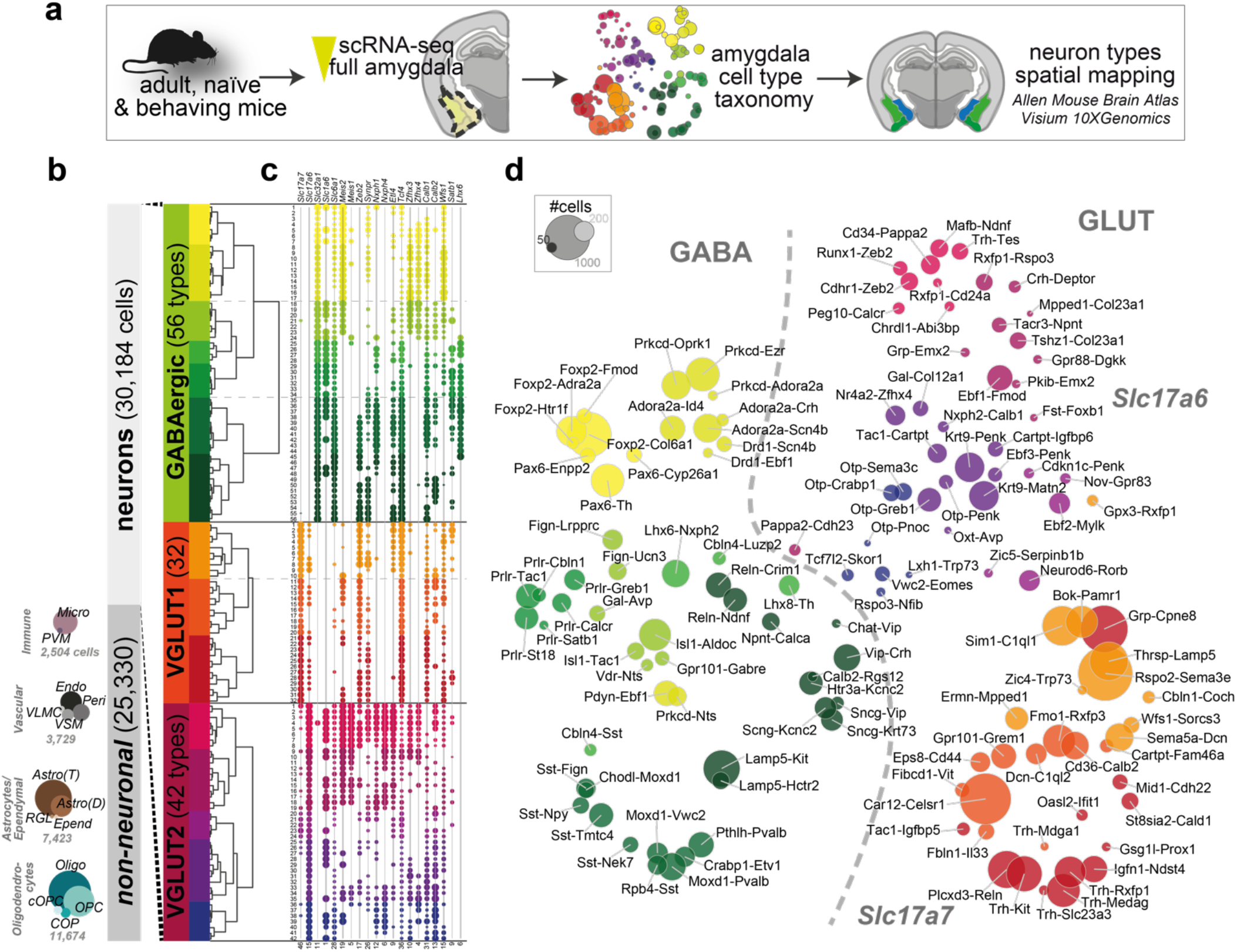
Amygdala neuron taxonomy. **a** Experimental overview; single-cell RNA-seq of whole amygdala of 23 mice to construct the cell type taxonomy, spatially mapped to distinct amygdala nuclei. **b** Data structure: of 55,514 whole-cell transcriptomes, 30,184 were high-quality neurons. Neuron clusters organized by neurotransmitter type (GABAergic (13,006 cells, 56 clusters) or glutamatergic (17,178 cells)), and glutamatergic cells expressed either vesicle transporter *Slc17a6* (VGLUT2, 5,231 cells, 42 clusters) or *Slc17a7* (VGLUT1, 11,947 cells, 32 clusters). Non-neuronal cell types indicated by circles; area corresponds to cluster size. **c** Cluster-wise gene expression of broad, modular markers; circle size represents cluster 75th percentile UMI count; maximum indicated at the bottom. **d** tSNE visualization of all 130 neuronal clusters by cluster average, colored by clade; circle area corresponds to cluster size.

To identify robust but refined molecular cell types, we performed clustering separately on GABA (13,006 cells), VGLUT1 (11,947 cells), and VGLUT2 (5,232 cells) (see Methods), and built a detailed neuronal cell type taxonomy of the mouse amygdala. GABAergic cells were clustered into 56 distinct cell types, VGLUT1 cells into 32 types, and VGLUT2 cells into 42 types (Figure 1b-d). Clusters were defined by their modulatory and/or exclusive expression of a multitude of enriched genes (Suppl. Table 2). We chose two marker genes for a naming scheme to describe each cell type as precisely as possible. For ease of handling and visualizing the many cell types, we defined more highly correlated clusters as clades (depicted by a color scheme, Figure 1b, d) for each of the three main groups: 7 clades for 56 GABAergic, 3 clades for 32 VGLUT1, and 5 clades for 42 VGLUT2 clusters. We detected all cell types in all sampled mice, both naïve and fear-conditioned, at ratios expected according to the sampling depth (Extended Data Fig. 1), meaning that CFC did not affect the cell type identity of any neuronal population.

### Spatial distributions reflect molecular neuron class

The amygdala is a collection of anatomically and functionally defined nuclei belonging to three main structural-developmental compartments: striatal, cortical, and subplate amygdala nuclei. To reconstruct the molecular cell types’ spatial context, we first plotted the expected localization of the three main classes (GABA, VGLUT1, VGLUT2) from expression densities per amygdala nuclei, in the Allen Mouse Brain Atlas *in situ* hybridizations^25^, for *Gad2, Slc17a7*, and *Slc17a6*, respectively (Figure 2a). As expected, *Gad2* was enriched in the striatal nuclei, especially the central and medial amygdala (CeA and MeA), and sparse in all other amygdala nuclei. Glutamatergic markers were entirely absent from CeA. In the other amygdala nuclei, VGLUT1 and VGLUT2 showed little overlap, where *Slc17a6* was striatal and cortical (MeA and CoA-anterior), and *Slc17a7* more restricted to cortical subplate nuclei (LA, BLA). This refines our previous report of VGLUT1 and VGLUT2 subtypes dividing largely according to the telencephalon-diencephalon boundary^17^, within the ventral compartments of the telencephalon.

**Figure 2.**
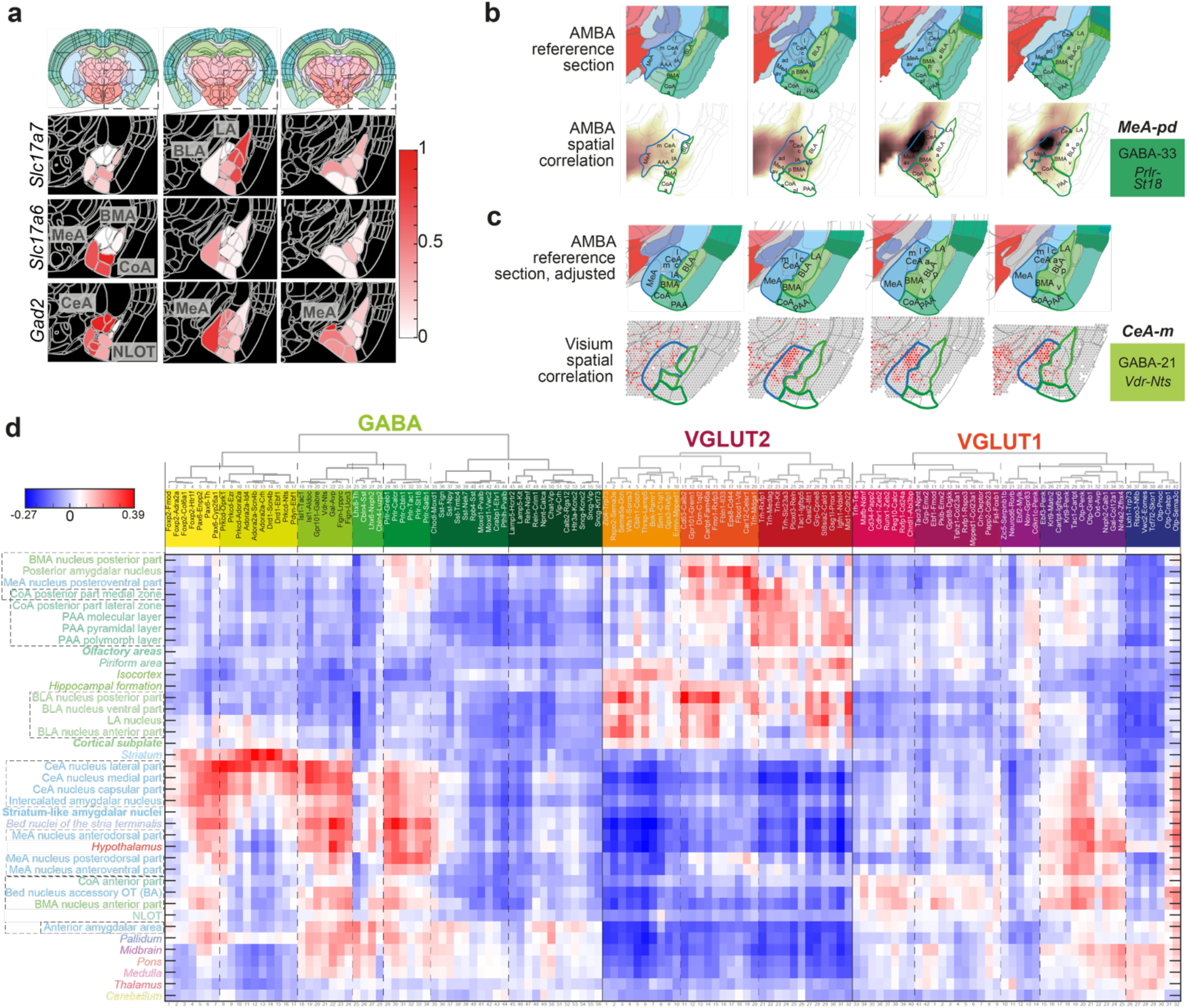
Spatial distribution of neuronal types in the amygdala. **a** The main splits of amygdala taxonomy; neurotransmitter and glutamate transporter expression are regionally distinct along the amygdala anterior-posterior (a-p) axis, and between amygdala nuclei. Expression values calculated and normalized per gene across three a-p coronal sections of the Allen Mouse Brain Atlas, and segmented nuclei are colored by relative expression levels. Top panel shows the anatomical context of the sections below (approximate; exact matches were not available), from the Allen Mouse Brain (AMB) Reference Atlas. **b-c** Examples of inferred spatial distribution of scRNA-seq clusters, by correlating transcriptomic profiles with AMB ISH Atlas (**b**) or Visium spatial transcriptomics (**c**). The top rows show amygdala zoom-ins on four a-p (left-to-right) AMBA reference sections, with amygdala nuclei labelled. Bottom rows visualize correlation with the indicated cluster. For every cluster and every voxel (AMBA) or spot (Visium), a correlation coefficient heatmap is shown (**b** dark, high; white, low; **c** red, high; grey, low). The top-scoring amygdala region is highlighted in bold. **d** Spatial correlation of all amygdala neuron clusters with the dissected amygdala nuclei, related or neighboring regions, and other higher-order grey matter structures. Regions are sorted by correlation similarity. Region names are colored according to the AMBA convention; regions that were not (fully) dissected are in italic; higher-order amygdala regions are in bold.

To guide our analysis of the 130 neuronal types, we next inferred spatial distributions for each cell type by correlating scRNA-seq expression profiles with Allen Mouse Brain Atlas (AMBA) single-gene ISH-based volumetric maps^17^ (see Methods). Despite the comparatively coarse resolution of 200μm voxels of the ISH-atlas, this helped place most cell types within amygdala nuclei (Fig. 2d, Suppl. Table 3). For smoother visualization, we parcellated volumetric ISH data into 25μm voxels and performed linear interpolation (Fig. 2b and subsequent AMBA spatial correlation figures). We found that spatial correlation often followed the molecular taxonomy of cell types, that is, cell types related by gene expression also shared regional origin, with some interesting exceptions (Fig. 2, Extended Data Fig. 2). For example, VGLUT1 clusters largely kept their organization and split according to correlation with BLA vs. CoA, mixing with only a handful of VGLUT2 clusters that localized to the LA. A separate branch of clusters, entirely from VGLUT2 neurons, correlated with anterior CoA. In the MeA and BMA, neurons from VGLUT2 and GABA classes intermixed.

Spatial correlation analysis also revealed the relations of amygdala cell types to expression profiles found elsewhere in the brain. As expected, CeA and IA clusters were molecularly related to their dorsal neighbor, caudoputamen. By contrast, MeA nuclei were similar to each other, the bed nuclei of stria terminalis (BST), and the hypothalamus—all known to share functions and circuitries in (sexually dimorphic) social behaviors. Both subplate and cortical amygdala areas were related in their high correlation with distinct glutamatergic populations, and as such resembled expression profiles also found in the isocortex and hippocampal formation.

Not all neuronal cell types showed high – or any – correlation within amygdala regions. We manually validated these by inspecting single marker genes (AMBA ISH, GENSAT), the literature, and other datasets, and found two explanations: (a) accidental sampling of neighboring structures (e.g., GABA 12, 14 and 15, caudoputamen; VGLUT1 16 and 17, ventral CA3 and DG; GABA 25, globus pallidus); and (b) spatial correlation reached its detection limit. In the latter case, cell types were either restricted to a very small area (e.g., *Avp-Gal* GABA 22, CeA, Fig. 3f; and *Oxt-Avp* VGLUT2 32, MeA), rare (e.g., olfactory clusters VGLUT2 1-17, Extended Data Fig. 7), or lacked spatial patterning (e.g., GABAergic interneurons GABA 35-56, Extended Data Fig. 4).

**Figure 3.**
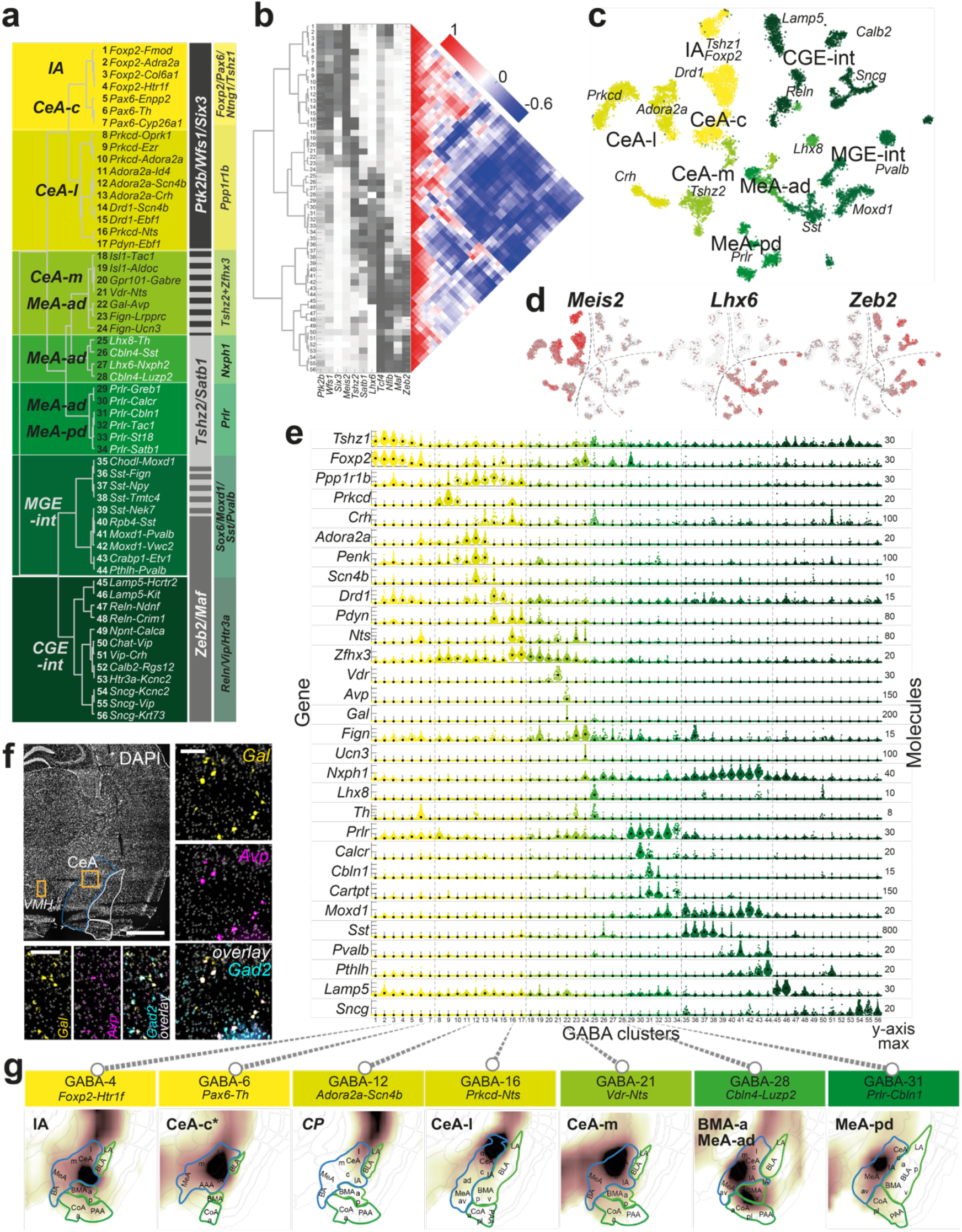
GABAergic cell types of the mouse amygdala. **a** Dendrogram of GABAergic cell types, with cell type number and two-gene identifier. Modular genes defining branching points across the dendrogram (grey), and in the specific clades (yellow-green) are highlighted on the right; likely location on the left (based on AMBA spatial correlation). IA, intercalated amygdala; CeA, central amygdala; -c, capsular; -l, lateral; -m, medial; MeA, medial amygdala; -ad, anterodorsal; -pd, posterodorsal; CGE, caudal ganglionic eminences; MGE, medial ganglionic eminences; int, interneurons. **b** GABA-types dendrogram with correlation matrix (right) and heat map of branch point marker gene expression (center). **c** tSNE visualization of all 13,006 GABAergic cells, colored by clade, labelled by main cross-cluster marker genes, and likely location. **d** tSNE (as in **c**) colored by expression of indicated branch point marker genes; white, 0; grey, low; red, high. Dashed lines indicate territories of the three main patterning gene modules. **e** Genes enriched to single or groups of clusters, visualized as single-cell violin plots. Each dot represents one cell; black square, median cluster expression, number of molecules (y-scale maximum) on the right. **f** Multiplexed fluorescent *in situ* hybridization of *Gad1* (GABA), *Gal*, and *Avp* in a representative posterior section. Top left, section overview, scale bar, 500μm. Right, CeA; bottom, VMH (Hyp) both scale bar 100μm. **g** Examples of inferred spatial distributions of seven distinct cell types, visualized in a single relevant section with high correlation to AMBA.

For independent spatial validation, we performed spatial transcriptomics (Visium, 10xGenomics) on 8 coronal sections along the anterior-posterior axis of the amygdala. This approach did not match AMBA-correlation in 3D resolution but detected and quantified ~3,000 genes simultaneously, per data point (mRNA-capture spot). As for AMBA volumetric data, we inferred the spatial distributions of the 130 scRNA-seq neuron types in this dataset (Fig. 2c, Extended Data Fig. 3, Suppl. Table 4). This approach independently validated the generalized AMBA spatial correlation patterns for most clusters. For example, VGLUT1 types were enriched in the basolateral nuclei, whereas VGLUT2 were detected in both cortical and striatal nuclei, with the exception of central amygdala that had only GABA types. Inhibitory interneurons lacked spatial enrichment.

### Organizational principles behind projection vs. local GABA neurons

Our analysis revealed 56 GABAergic neuron types covering the unique combination of cortical and subcortical amygdala compartments, where GABAergic cells are local inhibitory interneurons (e.g., *Sst+, Lamp5+*) and striatal compartments, where projecting inhibitory neurons, called medium spiny neurons (MSNs, e.g. *Ppp1r1b+*), dominate (Fig. 3 a, c, e). To understand without bias how this organization was reflected in expression patterns, we first analyzed which genes were most decisive in determining the split at each level of the cluster dendrogram (branch point marker genes, see Methods). The resulting list of genes revealed hierarchical and combinatorial expression of many (homeodomain) transcription factors, consistent with their role in specifying and maintaining neuronal identity.

Determining the first two dendrogram splits were *Wfs1*, *Meis2*, and *Ptk2b* vs. *Maf* and *Zeb2*, enriched on either distal ends of the tree (Fig. 3a, b, d). Indeed, this organization indicated a split between projecting vs. local interneuron populations: projecting markers *Ptk2b* (telencephalon^17^) and *Zfhx3, Zfhx4, Meis2* (spinal cord^26^) overlapped with striatal MSNs-like clusters. Local/interneuron markers *Nfib*, *Nfix*, *Tcf4, Satb1*, and *Prox1*^26^ were expressed among canonical interneuron types, with the notable exception of *Foxp2*-clusters (GABA 1-4) also expressing *Nfib, Nfix*, and *Tcf4* (Fig. 3b). *Tshz2, Satb1*, and *Lhx6* defined cell types in the center of the tree, with some overlaps with both distal marker sets. The expression of *Lhx6* suggested that these central clusters were medial amygdala GABAergic types^27^.

Consistent with these observations, spatial correlation analysis of GABA clusters (Fig. 2d, 3g) revealed the following generalized pattern: projecting *Ptk2b*/*Wfs1*/*Meis2* clusters were in the IA and CeA, and *Tshz2/Satb1/Lhx6* clusters were strongly enriched in the MeA. Cell types expressing a local interneuron *Maf*/*Zeb2* signature were generally found in all compartments, although they often showed low or no correlation with specific amygdala regions in AMBA volumetric data, discussed below (Suppl. Table 3).

### Inhibitory neurons of valence-learning modulation and output

Inhibitory cell types GABA 1-24 expressed known MSN markers (e.g., *Penk, Pax6, Gpr88, Ppp1r1b*), also described in scRNA-seq studies of other striatal regions^17,28–31^. They correlated strongly with striatal amygdala nuclei IA and CeA (Fig. 2c, Extended Data Fig. 5, Suppl. Table 3), two regions characterized in detail for their involvement in fear conditioning, and more generally, aversive and appetitive behaviors. We found that several MSN types followed the classical D1/D2 convention based on their dopamine receptor expression (here, *Drd1*, D1; *Adora2a*, D2), however, this was neither an exclusive nor a mutually exclusive hallmark of these neurons.

*IA (GABA 1-4):* First, four *Foxp2-Tshz1* types (GABA 1-4) were located to the intercalated amygdala nuclei (IA), embedded between the BLA-CeA regions. *Foxp2* is a known marker for IA intercalated cells (ITCs)^32^, which receive input from the BLA and modulate CeA activity^33^. *Tshz1* was described in a subgroup of D1-MSNs in the nucleus accumbens (NAc)^29^, and patch-specific MSNs we found in the dorsal and ventral striatum^17^ revealing an interesting relationship between ITCs and other striatal MSNs. The *Foxp2*-types all expressed *Drd1, Myh7*, the serotonin receptor *Htr1f*, and transcription factors *Pax6 and Pbx3*. Between them, ITCs were molecularly distinct, based on several genes: *Fmod*, sodium voltage-gated channel *Scn10a*, and adrenoreceptor *Adra2a* or *Col6a1*. We found no preferential localization of the molecular *Foxp2* clusters to the dorsal or ventral IA, as was recently reported for two functionally distinct ITCs^13^.

The CeA is a major output region of the amygdala. Relative to its size, it contributed a high number of inhibitory neuron types, spanning clusters GABA 5-23 (33% of GABA cells). Most of these cells expressed MSN markers enkephalin (*Penk), Six3*, and *Ano3*, as well as axon guidance-encoding gene *Epha4*, and zinc finger homeobox TF-encoding *Zfhx4* and *Zfhx3*.

*CeA-c (GABA 5-7):* Three cell types related to ITCs, GABA5-7, had lower levels of *Foxp2*, *Tshz1*, and *Pbx3*; they expressed the tachykinin receptor *Tacr3*, specifically *Tshz2* and *Enpp2*, *Nts* and *Th*, or *Cyp26a1*. Their location was in the neighboring capsular central amygdala (CeA-c).

*CeA-l/CP (GABA 8-17)*: Inhibitory cell types GABA 8-17 expressed higher levels of *Ppp1r1b*, encoding the dopamine-dependent central regulatory protein DARPP-32^34^ and *Crym*, described as a marker for MSNs in the medial and caudal striatum^30,35^. Consistent with this, *Ppp1r1b* clusters showed strong spatial expression correlation with the lateral CeA and neighboring CP (GABA 12, 14, 15). We found that the transcription factor *Zfhx3* largely distinguished CeA-MSNs (*Zfhx+)* from CP-MSNs *(Zfhx3-)* (Fig. 3g, Extended Data Fig. 5). Subgroups of inhibitory CeA-l/CP neurons expressed TF *Foxp1* (GABA 11-15) and D1-type MSN marker *Drd1* (dopamine receptor 1, GABA 14-15), D2 marker *Adora2a* (GABA 10, 13), or both (GABA 11-12). The *Ppp1r1b* clade also included well-described neurons specific to the CeA and positive for protein kinase C delta *Prkcd* (GABA 8-10). *Prkcd* was expressed in combination with kappa-type opioid receptor *Oprk1*(and *Dlk1, Cyp26b1)*, *Ezr*, or *Adora2a*. These neurons likely constitute one or several types described for their involvement in both fear and appetitive behaviors^3,36,37^. The smallest of the *Prkcd* clusters, GABA10, coexpressed *Adora2a* and the calcitonin receptor-like *Calcrl*, which were previously shown in a D2-MSN-like neuron type of the capsular CeA (CeA-c)^3^. Corticotropin releasing hormone *Crh* (also known as CRF) was expressed in D1 and D2 *Ppp1r1b* CeA-l types GABA 12-17, likely marking a separate CeA-l neuronal population extensively studied in anxiogenic circuits, fear response type and memory^38–40^. Several *Crh*-types also expressed prodynorphin *Pdyn* and *Isl1*, for instance with *Scn4b* (GABA 14), or *Ebf1, Unc13c*, and *Syndig1l* (GABA 17). In contrast with previous reports^36,41^, one D1-type (GABA 16) coexpressed *Prkcd* and *Crh*. This type was further marked by *Vipr2* and *Nts*. CeA-l type GABA 17 also expressed somatostatin *Sst* and may constitute the CeA-l type reported to antagonize *Crh*-CeA-l neurons in flight-versus-freeze fear behavior^38^.

### Neuropeptide-rich GABA classes in the medial CeA and across the MeA

Transcription factors *Tshz2* and/or *Zfhx3* marked the GABAergic populations 18-24, which we found to be most strongly correlated with the medial CeA (CeA-m) and the anterodorsal medial amygdala (MeA-ad). Several types (GABA 18-20) distinctly expressed the orphan G-protein coupled receptor *Gpr101*, for example, in combination with *Pdyn and Isl1* and *Tac1/Dlk1/Dgkk/Asb4* or *Aldoc*. *Gpr101*-expressing MSNs were previously found enriched in matrix-MSNs (D1 and D2) across the striatum^42^. One interesting, distinct population among CeA-m neurons, GABA 21, highly specifically expressed vitamin D receptor *Vdr*. *Vdr* expression was recently reported in the brain of Vdr-CRE reporter mice, especially in the cortex, caudate putamen, reticular thalamic nucleus, and amygdala^43^. Another type, GABA 22, coexpressed neuropeptides galanin and vasopressin (*Gal, Avp*), and was enriched in structures outside the amygdala, such as hypothalamus and bed nuclei. We used RNAScope *in situ* hybridization to validate a small population located to the MeA-ad (Fig. 3f), consistent with^44^. Finally, two types localized to the MeA expressed *Fign* and *Nts*, and either *Lrpprc* and *Th* (GABA 23) or the peptide hormone precursor *Ucn3*, implied in social behavior^45^, and tachykinin *Tac1* (GABA 24). GABA 25 had a unique gene signature of *Gpc5, Gbx1, Lhx8, Tacr3, Megf11, Th*, and *Fibcd*, several of which located it to the neighboring pallidum (TELINH1^17^). GABA 26-28 expressed GABAergic synapse organizer *Cbln4* and (predicted) neuropeptides *Sst*, *Nxph2*, or *Luzp2*. Most inhibitory cell types that we localized to the anterior amygdala area (AAA) and medial amygdala expressed *Lhx6*. This was consistent with previous reports of *Lhx6+* MeA-projecting inhibitory cell types^27^.

In the dorsal medial amygdala, we found six distinct GABAergic cell types (29-34) expressing receptors for prolactin (*Prlr)*, gonadal steroids androgen and estrogen (*Ar, Esr2)*, and estrogen responsive *Greb1*. Allen Mouse Brain *in situ* correlation indicated a gradient of these *Prlr*-types along the anterior-posterior axis of the MeA. Much of their molecular heterogeneity could be described by specific markers, such as calcitonin receptor *Calcr* (anterior), estrogen receptor *Esr2*, *Dkk3/Tac1/Cartpt*, *Pappa, St18*, or *Moxd1* (posterior). *Prlr* in the MeA and related circuits has been studied with respect to maternal behavior^46,47^. Therefore, and based on gonadal steroid receptor expression, the MeA-*Prlr* populations likely constitute neurons described in the pheromone-processing pathways to the medial preoptic area, activated selectively in males and virgin females, but not mothers^48,49^.

### Inhibitory interneurons are heterogenous, scarce, and dispersed

The remaining types GABA35-56 were largely marked by *Maf/Zeb2*, with some overlap with CeA-m/MeA-markers *Satb1/Tshz2*. Among their most enriched genes were canonical cortical interneuron markers, such as *Sst, Pvalb, Vip*, or *Sncg*. In agreement with a plethora of research and recent scRNA-seq studies^50–52^, the main division between the types was developmental, i.e., whether they derived from the medial or caudal ganglionic eminences: MGE types included canonical *Sst* and *Pvalb* interneurons, and CGE types expressed *Lamp5, Reln, Vip*, or *Sncg*.

Most MGE populations (GABA35-44) expressed voltage-gated potassium channel KV3.2 gene *Kcnc2*, coherent with fast-spiking *Sst* and *Pvalb* phenotypes. *Satb1* marked *Sst* neurons, while *Igfbp4* was higher in *Moxd1, Pvalb*, and *Pthlh* cells. Among CGE-types (GABA45-56), we found *Lamp5/Hcrtr2* (including *Kit*), *Reln, Chat*, and *Vip/*Crh interneurons. *Kcnc2* was expressed in a subgroup, marked by *Npas1, Rgs12, Htr3a*, and *Sncg. Pvalb* interneurons came in two types: a *Pthlh/Unc5b-Pvalb* population and a *Moxd1/Gpr83-Pvalb* type, both consistent with dorsal striatum *Pvalb* interneurons^28^. We found a small population of cholinergic neurons expressing high levels of *Chat* (GABA50), but several, including *Vip-Crh* (GABA51), expressed low levels of choline transporter *Slc5a7*.

Canonical interneuron types have been studied in the amygdala. They are scarce, however. For example, in the mouse BLA, GABAergic interneurons make up only an estimated 5-9% of neurons between the dominant glutamatergic principal cells^11^. Accordingly, spatial correlation analysis detected no local enrichment to specific amygdala compartments (Fig. 2d, Extended Data Fig. 3). Genes that we found to mark only certain interneuron types and no other amygdala neurons were expressed across the amygdala in the Allen Mouse Brain ISH Atlas. For example, *Pthlh* or *Vip* positive cells were extremely scarce and dispersed across several subplate and cortical amygdala nuclei (Extended Data Fig. 4).

### Excitatory cells class by anatomical compartment

Among glutamatergic cells, we found that the division between VGLUT1 and VGLUT2 neurons consistently marked the first split in transcriptome-wide correlation. Beyond the vesicular glutamate transporters *Slc17a6* or *Slc17a7*, other genes distinguished them (Fig. 4c, e): vesicular transporters *Slc1a6* (glutamate/aspartate), *Slc30a3* (zinc) and *Slc6a1* (GABA), transcription factors *Tcf4, Pbx3*, and others (e.g., *Baiap3, Prdm8, Nov*). Therefore, clusters coexpressing *Slc17a6* and *Slc17a7*, at times even at comparable levels, were nevertheless unequivocally assigned to one or the other group. VGLUT2 cell types were also the more molecularly diverse class: from only 5,231 VGLUT2 cells we identified 42 distinct clusters, while 11,947 VGLUT1 cells yielded 32 distinct clusters. As predicted by AMBA *in situ* of *Slc17a7* and *Slc17a6* alone (Fig. 2a), clusters belonging to either group showed spatial enrichment along the anterior-posterior axis, and to amygdala compartments; VGLUT1 posterior and in BLA, PAA, PA, and CoA-p and VGLUT2 anterior and in BMA, CoA-a, BA, and MeA (Fig 2d, 4a). *Slc17a6/Slc17a7* double positive types were also restricted to distinct regions (Fig. 4 f-g), such as the LA, CoA, and an olfactory area associated with cortical amygdala, NLOT, discussed below.

**Figure 4.**
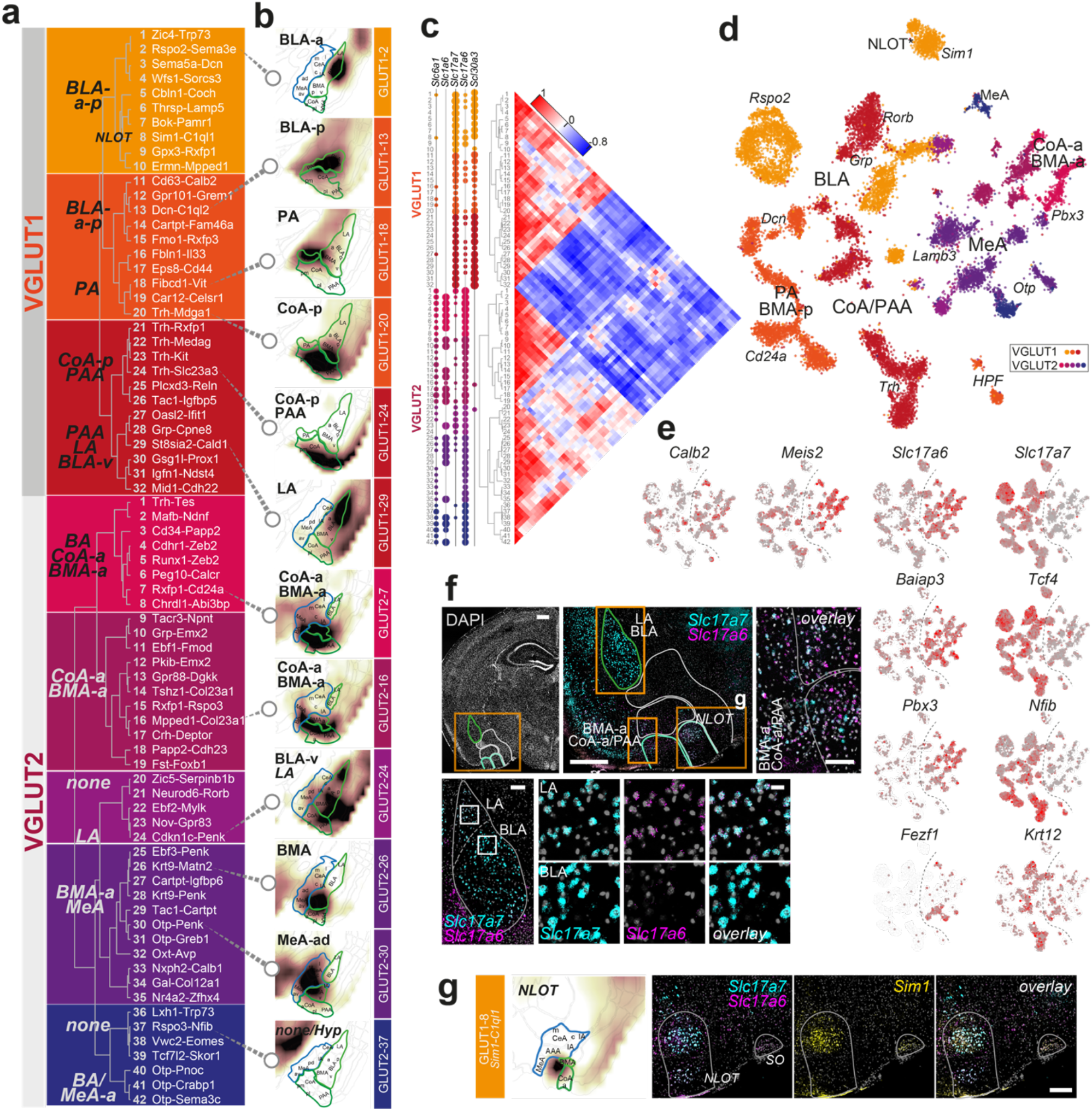
Glutamatergic diversity. **a** Dendrogram of glutamatergic cell types of the VGLUT1 and VGLUT2 classes, with cluster number and two-gene identifier; likely location (based on AMBA spatial correlation) indicated on the left and **b** examples of inferred spatial distributions of twelve distinct cell types, visualized in a single relevant section with high correlation to AMBA. BLA, basolateral amygdala; NLOT, nucleus of the lateral olfactory tract; PA, posterior amygdala; CoA, cortical amygdala; PAA, piriform amygdalar area; LA, lateral amygdala; BMA, basomedial amygdala; MeA, medial amygdala; a, anterior; p, posterior. **c** VGLUT1/VGLUT2 types dendrograms with relevant transporter gene expression, left, and correlation matrix, right. **d** tSNE visualization of all 17,178 glutamatergic cells, colored by clade, labelled by main cross-cluster marker genes, and likely location. **e** tSNE (as in **d**) colored by expression of indicated branch point marker genes; white, 0; grey, low; red, high. Dashed line marks VGLUT1-VGLUT2 class border. **f** Multiplexed fluorescent *in situ* hybridization of *Slc17a7* (VGLUT1) and *Slc17a6* (VGLUT2) highlighted in LA, BLA (scale bar, 20μm), and anterior CoA/BMA (scale bar, 100μm), in a single representative anterior section. Top panel section overview (scale bars, 500μm); LA/BLA overview (scale bar, 100μm). **g** Left, inferred spatial distribution of cluster GLUT1-18. Right, multiplex FISH validating *Slc17a7+Slc17a6+Sim1+* cells in the NLOT. SO, supraoptic nucleus. Scale bar, 100μm.

### The famous, and the unknown: Basolateral principal cells

Basolateral amygdala glutamatergic cells (BLA principal cells) were previously described in depth by their topographical distribution, circuitry (projections), and differential valance assignment, that is, their encoding of positive versus negative valance^2,4,53^. But their molecular description has to date been restricted to a handful of markers, such as *Rspo2, Ppp1r1b*^2^, and more recently, *Fezf2*^4^. Consistent with the diverse functional properties and topography of glutamatergic cell types that mapped to the BLA, we found great molecular diversity among them. These cell types showed distinct enrichment or gradients along the a-p axis, and we present genes that precisely mark their identity (Extended Data Fig. 6, Fig. 5a, b). They were almost exclusively VGLUT1, although several co-expressed low levels of *Slc17a6* (Fig. 5).The most cell-rich branch, VGLUT1 1-4, expressed *Man1a/Dkk3* and was localized in an a-p gradient across the BLA. Of these, the largest cluster (VGLUT1 2), localized to the anterior BLA, expressed low levels of known BLA marker *Rspo2*, and specifically, *Satb1, Neurod6, Rph3a*, and *Cyp26b1*. Two types (VGLUT1 3-4) stretched across the full BLA, and were marked by *Sema5a, Dcn, Tgfb2* (3) or *Wfs1, Sorcs2*, and *Bdnf*. Six more posterior types (VGLUT1 10-15) expressed combinations of *Meis2, Calb2*, and *Dcn*, and *Mpped1/Tac1, Cd36, Gpr101, Cartpt* or sialyltransferase *St8sia2*, and *Ptpru*. All but the two most posterior types (VGLUT1 12-13) expressed serotonin receptor *Htr2c*.

**Figure 5.**
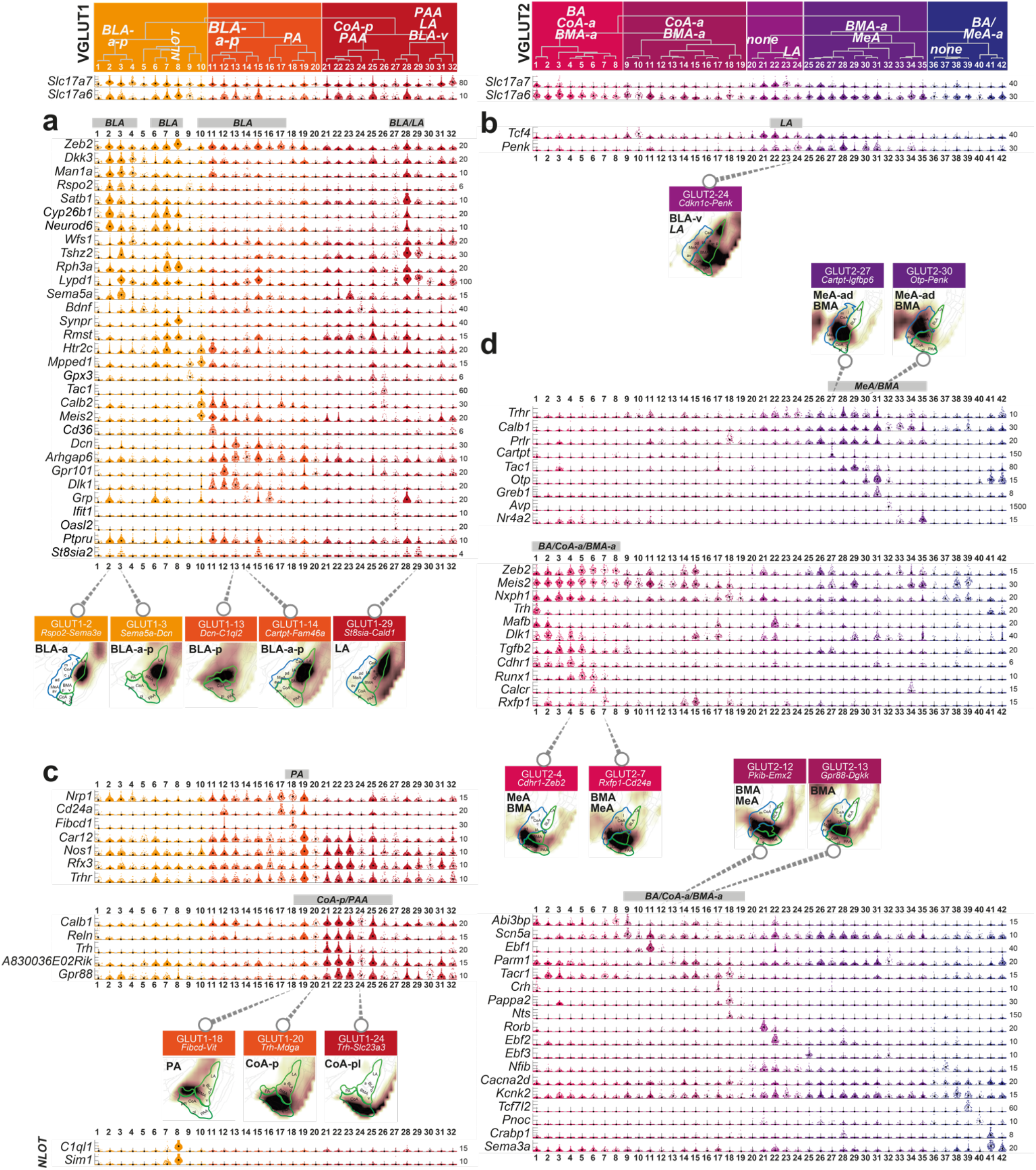
Highly diverse glutamatergic cell types of the amygdala as part of the **a-b** subcortical (BLA/LA) or **c-d** olfactory structures. Genes enriched to single or groups of cell types, visualized as single-cell violin plots. Each dot represents one cell; black square, median cluster expression, number of molecules (y-scale maximum) on the right. Examples of inferred spatial distributions of distinct cell types, visualized in a single relevant section with high correlation to AMBA, below each expression panel.

We identified three VGLUT1 types (27-29) enriched to the most dorsal aspect of the BLA, the lateral nucleus (LA). The most prominent among them (VGLUT1 28) was distinctly marked by the neuropeptide gastrin releasing peptide *(Grp). Grp*-expressing pyramidal neurons were previously identified in the LA, where a local circuitry with *Grpr*-expressing LA interneurons influenced certain aspects of fear memory^12,54^. Indeed, we found sparse, but specific expression of *Grpr* in several GABAergic interneurons, such as *Vip*-expressing GABA 50 and 51, and *Pvalb*-type GABA 41, and a single glutamatergic type, *Trh*-expressing VGLUT1 21 of the CoA. Interestingly, LA-*Grp* cells shared expression of *Cyp26b1, Neurod6, Satb1*, and others almost exclusively with the BLA-*Rspo2* type (VGLUT1 2).

Of note, a single VGLUT2 type (24) showed a similar spatial distribution as the VGLUT1-LA-types, in particular, VGLUT1 27. Both were enriched not only in the LA but also to the ventral nucleus of the BLA. VGLUT1 27 curiously expressed *Oasl2* and *Ifit1*, both associated with interferon response. VGLUT2 24 had a distinct expression profile, such as *Stxbp6, Id2*, and *Penk* (Fig. 5b). Their similar spatial alignment was thus very likely true, not based on coincidental expression of odd markers.

### Glutamatergic cells of the olfactory amygdalar areas

The majority of glutamatergic cells outside the subplate nuclei came from the olfactory-associated amygdala areas. Here, we identified three main branches, largely reflecting different components of the accessory and main olfactory systems.

We found *Calb1*^+^*Zeb2*^−^ best described putative accessory olfactory system in both VGLUT1 and VGLUT2 neurons (Fig. 5c, d). For example, VGLUT1 types 20-26 localized to the posterior CoA and piriform amygdalar area (PAA), and several showed great molecular similarities with external piriform areas. Most were also marked by expression of *Trh*, *Reln*, and *A830036E02Rik* (Fig. 5c). *Trh+* amygdala neurons were described in anxiolytic behavior^55^, and recently, in suppression of male mating behavior through a CoApm-MeA connection^14^. In the latter study, CoApm-*Trh* cells projected to *Trhr*-expressing VGLUT2 cells in the anterior MeA. Although *Trhr* was more highly expressed elsewhere (e.g., in the posterior amygdala (PA), VGLUT1 19), *Trhr*-MeA cells correspond to clusters VGLUT2 27-35, also broadly marked by *Calb1*, specifically, by neuropeptides *Cartpt, Tac1, Avp/Oxt*, TFs *Otp, Nr4a2*, or estrogen-responsive *Greb1*. (Fig. 5d). Other VGLUT1 clusters lacking *Zeb2* were piriform (amygdala) area clusters VGLUT1 5, 9, and 30-32.

By contrast, anterior CoA clusters, VGLUT2 2-8, *did* express *Zeb2*. The anterior CoA is considered part of the main olfactory system. Other genes marking these anterior CoA clusters were *Tgfb2*, *Synpr* in combination with specific markers *Dlk1, Runx1, Calcr* or *Rxfp1* (Fig. 5d). These clusters were also present to varying degrees in the anteroventral MeA (MeA-av) and bed nucleus of the accessory olfactory tract (BA), and increasingly, the anterior basomedial amygdala (BMA-a). ISH data of single, precise marker genes, such as *Tgfb2*, confirmed this observation (Extended Data Fig. 7).

The next clade of putative main olfactory system VGLUT2 types 9-19 was increasingly distributed more dorso-laterally, to the surprisingly molecularly diverse BMA and anterodorsal MeA, or piriform amygdalar area (PAA). Like their ventro-medial neighbors, many still expressed *Meis2* and *Nxph4*, but *Synpr* and *Tgfb2* were not expressed. *Scn5a, Ebf1, Rxfp1*, and *Crh/Tacr1* were genes marking specific BMA subtypes. One distinct, small MeA cluster (VGLUT2 18) expressed *Pappa2/Prlr/Nts*. A subset of this population expressed *Cdh23*, which we found was dispersed across subcortical structures (Extended Data Fig. 7).

One large population of *Slc17a6/Slc17a7* double-positive neurons, VGLUT1 8, was marked by an array of distinct genes: *Sim1, C1ql1, Pou3f1*, and *Cbln2*. The cluster also shared the expression of more modular markers with other VGLUT1 clusters, such as *Zeb2, Rph3a*, and *Lypd1* (Fig. 5). Spatial correlation analysis, visual inspection of specific markers, and multiplexed *in situ* hybridization (Fig. 4f) placed this population to the nucleus of the lateral olfactory tract (NLOT), a cortical structure of the olfactory amygdala area embedded between the anterior striatal and classical cortical amygdala areas. Besides this population, we found that the neighboring CoA contained several more *Slc17a6/Slc17a7* double positive clusters, with VGLUT2 1 (*Trh-Tes*) a likely candidate in the anterior CoA, and *Trh*-VGLUT1 20-22 in the posterior CoA (Extended Data Fig. 7). Finally, two VGLUT1 clusters, 18 and 19, were from the posterior amygdala area (PA) and expressed *Pvrlr1* and *Fibcd1/Scrg1* (18) or *Car12, Trhr, Nos1*, and others (19). Although they are part of the accessory olfactory system, PA neurons were more closely related to their subplate neighbors.

### In silico trapping highlights CFC-activated cell types

Activation of neurons, for example during memory acquisition and recall, is associated with changes in gene expression. On the transcriptional level, upregulation of activity regulated genes, or immediate early genes (IEGs), can be detected. Activated cells within a neuronal population, based on upregulation of one or several IEGs, are termed the memory engram. The portion of activated neurons was shown to range from as few as 2-4% after CFC in the dentate gyrus to around half, in light-activated visual cortex^15,18,22^. We tested whether our approach could detect activated cells in any sampling timepoint after fear conditioning (context or cue), compared to the naïve (home cage, HC) control group (Fig. 6a). Comparing cells from the two groups, we found no difference in their median expression of IEGs. Instead, in many cell types, we found small subsets of cells that expressed IEGs at elevated levels (e.g. in the 90^th^ percentile over all neurons, Extended Data Fig. 8a). To test whether such subsets may constitute CFC-activated cell states within a cell type (the memory engram), we analyzed cells highly expressing IEGs *Arc, Bdnf, Btg2, Fos, Fosl2, Homer1, Npas4* or *Nr4a1*. Indeed, we found that activated subsets of cells were disproportionally from CFC, as compared to HC samples (Fig. 6b), and showed different time-dependent dynamics, both in fold change, and fraction of activated cells (Fig. 6d, Extended Data Fig. 8b).

**Figure 6.**
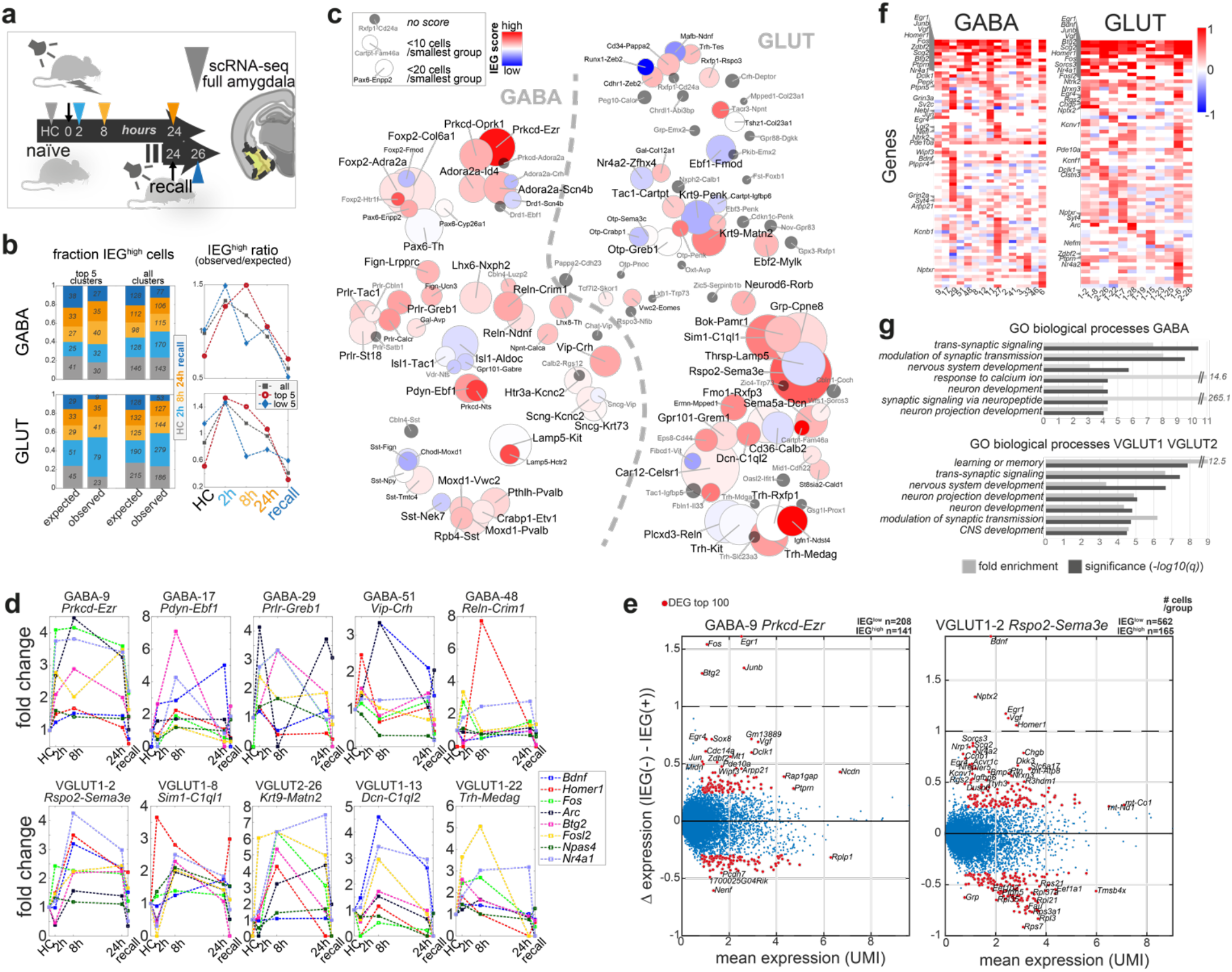
Cell type-resolved activity-regulated gene expression in cued fear conditioning. **a** Tone-cued fear conditioning, and scRNA-seq sampling timepoints after 2h, 8h, 24h, and 2h after next-day CS recall (no shock). HC, naïve home cage control. **b** Per timepoint; fraction and cell number (left) of cells highly expressing immediate early genes *Arc, Bdnf, Btg2, Fos, Fosl2, Homer1, Npas4* or *Nr4a1* (IEG^high^, 99^th^ percentile); “observed” vs. the fraction and number “expected” by chance, shown for the five top-responding, and all clusters. Right, resultant ratio (observed/expected), per timepoint. **c** Cluster-wise IEG score (see Methods), visualized as heatmap on tSNE (as Fig. 1d). Blue, low score; red, high score; grey, cluster too small to calculate score (<60 cells); small cluster label, smallest group size <20 cells (black), <10 cells (grey). Circle size represents cluster size. d Time-dynamic expression (fold change) of the eight IEGs, 90^th^ percentile, for the five top IEG-scoring clusters in among GABA (top) and GLUT (bottom) types. **e** Scatterplot of differentially expressed genes between activated (IEG^high^, 95^th^ percentile) and non-activated (IEG^low^) cells, for the highest IEG-scoring cell types among GABA and GLUT. Dots represent genes, red dots; top 100 DEGs with mean UMI >1. **f** DEGs frequently upregulated in activated (IEG^high^) cells (as e), per cluster. Blue, low; red, high. g Biological processes enriched in activated (IEG^high^) cells (as e), per class; FDR-adjusted q-values, and fold enrichment (DAVID).

Considering the amygdala’s highly heterogenous taxonomy, which cell types most responded to the CFC paradigm? Using the same panel of known IEGs, we summarized the fraction of activated cells, in CFC compared to HC controls, in an IEG score; calculated for each cell type (Fig. 6c, Extended Data Fig. 8c, Suppl. Table 5). Among the more deeply sampled cell types (≥20 neurons sampled per time point), the most strongly responding cell types in their respective class were the BLA *Rspo2-Sema3e* type (VGLUT1 2), and the CeA *Prkcd-Ezr* type (GABA 9), as measured by their IEG score. Other transcriptionally activated glutamatergic cell types were from the BLA (VGLUT1 13 *Dcn-C1ql2*, VGLUT1 3 *Sema5a-Dcn*, VGLUT1 12 *Gpr101-Grem1*), BMA (VGLUT2 26 *Krt9-Matn2*, VGLUT2 29 *Tac1-Cartpt*), nucleus of the olfactory tract (NLOT, VGLUT1 8 *Sim1-C1ql1*, VGLUT1 7 *Bok-Pamr1*), CoA (VGLUT1 22 *Trh-Medag*) and LA (VGLUT1 28 *Grp-Cpne8*). Among GABAergic neurons, CeA types GABA 17 *Pdyn-Ebf1* and GABA 8 *Prkcd-Oprk1*, MeA cluster GABA29 *Prlr-Greb1* and interneuron clusters GABA 51 *Vip-Crh* and 48 *Reln-Crim1* were most activated. Fig. 6d and Extended Data 8b show expression dynamics of 90^th^ percentile IEGs among these top activated, large cell type populations.

On the other hand, cell types least responding with an upregulation of classical IEGs in response to CFC included several clusters from the very same amygdala subregions, such as glutamatergic clusters of the BMA (VGLUT2 28 *Krt9-Penk*, VGLUT2 11*Ebf1-Fmod*) and BLA (VGLUT1 6 *Thrsp-Lamp5*, VGLUT1 11 *Cd36-Calb2*). Similarly, among GABAergic neurons, several CeA types did not deploy cells to the engram (GABA 19 *Isl1-Aldoc*, 18 *Isl1-Tac1*, GABA 6 *Pax6-Th*), and interneurons GABA 39 *Sst-Nek7* and 46 *Lamp5-Kit*, were similarly unresponsive.

Many smaller clusters (>60 cells total, <20 cells per time point; grey in Fig. 6c) likely also responded, although statistical confidence in their analysis is reduced. Among them, some notable examples of activated cell types were GABAergic intercalated cells (ITCs) GABA 4 *Foxp2-Htr1f*, compared to the non-activated ITC population GABA 1 *Foxp2-Fmod*. Several other CeA types also scored highly (GABA 16 *Prkcd-Nts*, GABA 5 *Pax6-Enpp2*), as did interneuron type GABA 45 *Lamp5-Hcrtr2*. Among the smaller glutamatergic populations, BLA type VGLUT1-14 *Cartpt-Fam46a* stood out. Further, both populations of the ventral hippocampus (vHPF), VGLUT1 16 *Fbln1-Il33* and VGLUT1 17 *Eps8-Cd44* were among the most activated cell types. This is in line with the vHPF’s involvement in emotional behaviors, including learned anxiety and fear memory^56–58^.

### CFC-activated neurons upregulate synaptic processes

Next, we analyzed which other genes covaried with the IEG panel: For each cell type, what distinguished activated from non-activated cells? We used the eight-IEG panel (*Arc, Bdnf, Btg2, Fos, Fosl2, Homer1, Npas4, Nr4a1*) to “trap” single cells expressing any IEG in the 95^th^ percentile over all neurons (activated cells). We then analyzed differential gene expression between this activated population (IEG^high^), vs. non-activated cells of the same cell type (IEG^low^) (Fig. 6e, Extended Data Figure 8, Suppl, Table 5). Across all cell types, besides the genes of the defined IEG panel (the “trap”), other known IEGs were also upregulated, such as *Egr1* (or Zif268), *Egr4, Junb or Scg2*. Globally, biological processes upregulated in association with the activated state were related to synaptic signaling and modulation of synaptic transmission/synaptic plasticity, but also development, projection morphogenesis, and learning and memory (Fig. 6g, Suppl. Table 5). For example, *Ntrk2* and *Vgf* are well-documented actors in synaptic plasticity and CFC acquisition^59,60^ that showed robust, activity-dependent transcriptional upregulation. *Ntrk2* (TrkB) encodes a receptor for neurotrophins BDNF and VGF, both of which we also found upregulated in activity-induced neurons. Activated TrkB is affects neurite outgrowth and synapse formation and plasticity via phosphorylation of CREB. We found many other robust candidates that were shared across activated cells in GABAergic and glutamatergic types, such as *Dclk1, Syt4, Clstn3, Pde10a, Ptprn and Zdbf2* (Suppl. Table 5).

Other upregulated genes were less widely-studied. Neuronal dense core vesicle component synaptotagmin 4 (*Syt4*) is involved in dendrite extension, and phosphatase *Ptprn* in vesicle-mediated secretory processes. Phosphodiesterase *Pde10a* regulates the intracellular concentration of cyclic nucleotides and was shown to be upregulated after LTP induction^61^. Pentraxin receptor *Nptxr* participates in synapse remodeling, and *Clstn3* is a regulator of synapse assembly^62^. Processes related to translation on the other hand, were downregulated in activated cells (via e.g. ribosomal protein genes) (Extended Date Fig. 8, Suppl. Table 5).

In addition, some class-specific differences were apparent (Fig 6g). For example, activated GABAergic cells specifically upregulated *Arpp21* that was shown to be essential in dendritic branching and complexity^63^, synaptic vesicle protein *Sv2c*, cytoskeleton-related transcripts *Tiparp* and *Wipf3*. GABA-specific activity-induced phosphatase *Ptpn5* (also known as STEP) regulates several effector molecules involved in synaptic plasticity, and *Plppr4* is important for axonal outgrowth during development and regenerative sprouting, and attenuates phospholipid-induced axon collapse. Although not exclusive, *Bdnf* was more widely induced in glutamatergic cells. Specific to activated glutamatergic cells were neuropeptide receptor *Sorcs3*, granin *Chgb*, and synaptic proteins *Nrxn3* and *Nptx2*, the latter implied in long-term plasticity. Voltage-gated potassium channels *Kcnv1* and *Kcnf1* were also enriched in activated glutamatergic cells.

Some cell type-specific patterns of activation also emerged: In activated cells of several GABAergic types of the medial amygdala (GABA-27, 29, 33) inhibitory synapse specifier *Lgi2*^64^ was highly expressed. *Egr4*, on the other hand, was restricted to activated cell types of the CeA (GABA-9, 11), BLA and LA (VGLUT1-2, 6,28).

### Single-cell expression correlation reveals CFC-associated learning gene modules

To test to what extend IEGs were co-expressed in individual cells, rather than just individually highly expressed (IEG^high^, Fig. 6), we next analyzed their pairwise correlation, per cluster, and experimental group (Fig. 7a). In analogy with IEG activation, IEG correlation was restricted to a subset out of all neuronal populations in the amygdala, and was consistently greater after CFC, compared to HC controls. Clusters with high IEG activation score often, but not always, had high IEG correlation, too.

**Figure 7.**
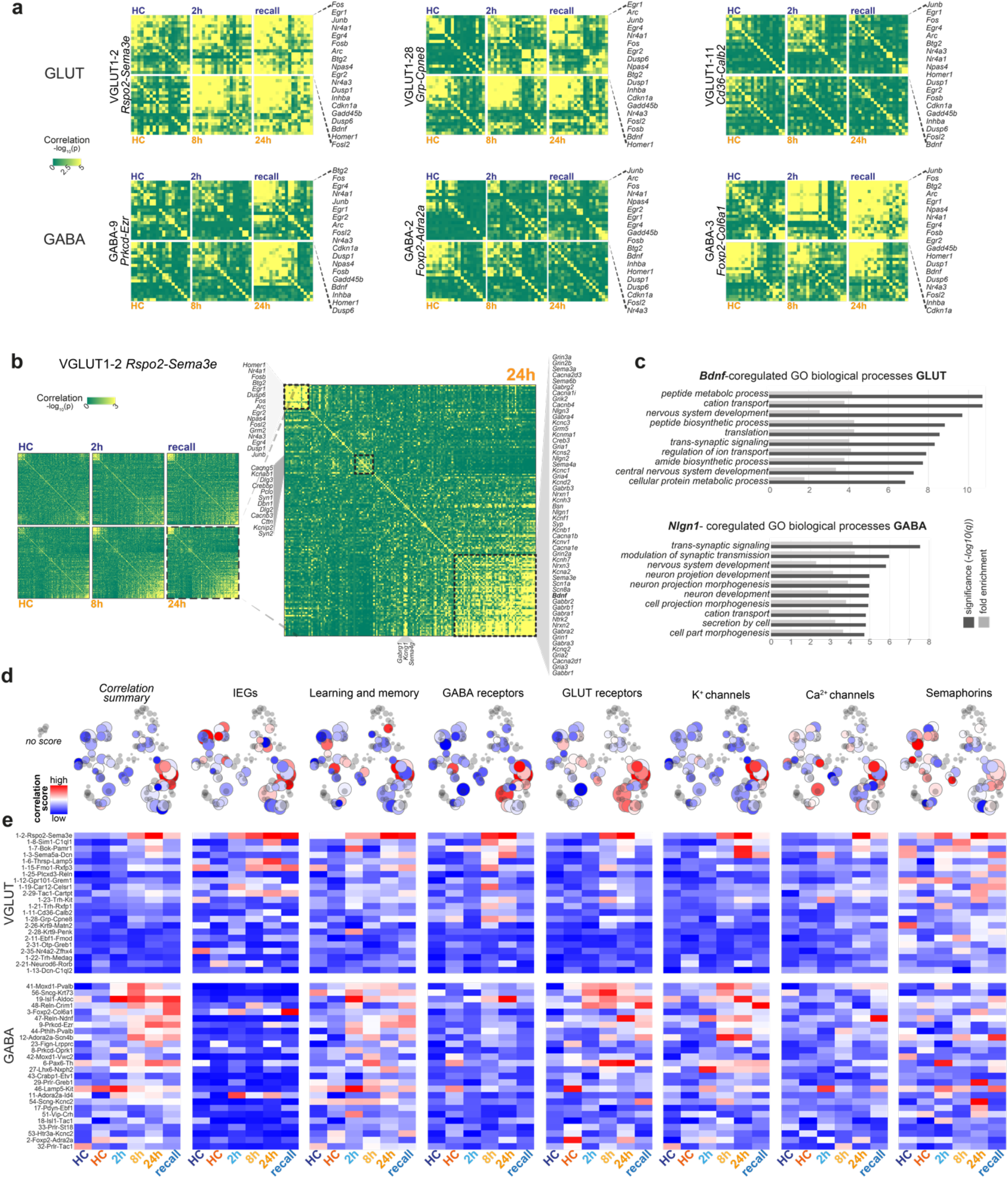
Gene expression correlation identifies cell types with CFC-activated learning gene modules. **a** Pearson pairwise correlation (green, low; yellow, high) of 18 IEGs (rows and columns) for six clusters, resolved by post-CFC sampling time point and batch-specific home cage (HC) control. For each cell type; top row, batch B (HC-2h-recall); bottom row, batch A (HC-8h-24h); genes are ordered by hierarchical clustering, indicated on the right. **b** Pairwise correlation of or 156 learning-related genes (rows and columns), in VGLUT1-2, per post-CFC sampling time point and batch-specific home cage (HC) control. Pearson coefficient (green, low; yellow, high), genes are ordered by hierarchical clustering. Zoom-in to 24h post-CFC (right), with three correlated gene expression modules highlighted.**c** Biological processes enriched among the top 200-genes correlated with *Bdnf* (GLUT) or *Nlgn1* (GABA); FDR-adjusted q-values, and fold enrichment (DAVID). **d** Cell-type resolved correlation, per gene module indicated, or all genes combined (“Correlation summary”), visualized as heatmap on tSNE (as Fig. 1b). Pairwise correlation score; blue, low; red, high; grey; no score (smallest group in cluster <20cells). Circle size represents cluster size. ***e*** Time-resolved correlation, per experimental group (post-CFC, HC controls). for large clusters (smallest group in cluster ≥20cells), resolved for sampling timepoints and batch-specific home cage (HC) controls, for gene modules as in ***d***. Pearson pairwise correlation score; blue, low; red, high.

For example, IEGs in the activity high-scoring GABAergic population *Prkcd-Ezr* were particularly highly correlated 24h after CFC, and 2h after recall. And while intercalated clusters *Foxp2-Adra2a* and *Foxp2-Col6a2* showed similar IEG activity scores (Fig. 6c), their correlation dynamics differed vastly; with only *Foxp2-Col6a2* strongly coregulating IEGs, particularly in the later sampling time points, and recall.

We further explored whether other known learning-related genes, receptors and channels displayed similar correlation dynamics (Fig. 7b, Suppl. Table 6). Indeed, we found increased correlation in a subset of clusters (Fig. 7c, Suppl. Table 6), and considerable differences between experimental groups (Fig. 7d). For example, VGLUT1-2 *Rspo2-Sema3e* showed strongly corelated, time-dynamic expression across all analyzed gene modules, while other BLA/LA clusters had more subtle responses (VGLUT1-11, 28), or lacked any CFC-induced coregulation (VGLUT1-13).

Among many analyzed genes, *Bdnf* (in GLUT) and *Nlgn1* (in GABA) were examples we frequently found highly correlated with both IEGs and other learning-related modules. To examine to what extent they may represent the wider orchestrated, transcriptional response to CFC, we analyzed each of their correlation, with transcriptome-wide gene expression (Extended Data Fig. 9, Suppl. Table 6). For both, correlation strength and frequency greatly increased after CFC, compared to HC controls. *Bdnf* most strongly correlated with co-TrkB ligand *Vgf*, and several of the same genes highly expressed in IEG^high^ cells (Fig. 6e-f); e.g., *Ptprn, Nptx2, Scg2, Lingo1* or *Scn1b*. *Nlgn1*-correlation among GABAergic clusters revealed several different candidates, such as synapse-acidifying proton pump *Atp6v0e2*, neurogranin *Nrgn*, and amyloid beta precursor *App*. Biological processes enriched concerned ion transport, transsynaptic signaling, development, and translation, peptide metabolism and biosynthesis (Fig. 7c).

### Coregulation characterizes cellular events after memory retrieval

We used IEGs to identify engram cells, since optogenetic and pharmacogenetic studies have shown that IEG-expressing neurons are required for memory recall. Transcriptional changes occurring in direct response to a memory recall event, however, have been described to a much lesser extent. For example, some, but not other reports, saw *Egr1* (Zif268) levels changed specifically after memory retrieval. In our data, *Egr1* was not specific to recall, but highly upregulated, with other IEGs, in all activated cell types. To the contrary, we found that the very IEGs that initiate processes required for memory acquisition were not elevated 2h after recall. Instead, IEG expression appeared *less active*: Both fraction of IEG^high^ cells, and fold change in their IEG expression, were lower after recall, than all other time points, including naïve HC controls, and the temporally close 24h-post CFC samples (Fig. 6b). Analysis of DEGs between same-batch recall and 2h only confirmed that cells expressed consistently higher levels of IEGs 2h post-CFC, than 2h after CS recall. Here, no other biological processes or pathways were significantly enriched for either condition (Suppl. Table. 5).

In our analysis of correlated genes, however, recall often showed highly coregulated expression, across gene modules and cell types (Fig. 7). Even IEGs, although lower expressed, were in many cases more correlated (i.e. more likely to be expressed in the same cell), 2h after recall, than 2h after CFC. In summary, more orchestrated gene expression, rather than elevated IEG expression, may describe cellular processes associated with memory consolidation and retrieval.

## Discussion

We presented a high-quality taxonomy of the full mouse amygdala, revealing molecular details over its functionally distinct and intertwined regions. Among regions studied in associative learning, such as fear or appetitive conditioning, BLA glutamatergic cells were more diverse, and more defined, than previously used markers revealed (e.g. *Fezf2, Rspo2*). Their molecular identities correlated with different anterior-posterior distributions, which could explain their role in encoding opposing valences. With regards to a more recent study performing scRNA-seq specifically on the BLA-LA^16^, we confirmed genes, such as *Cplx1* and *Lynx1* in BLA, and *Rorb* and *Myl4* in LA. Often expression was less BLA-specific in our dataset, which retained the context of all amygdala cells. For example, BLA marker *Rorb* was higher in CoA and *Cplx1* was also expressed across GABAergic interneurons. LA marker *Cdh13* and *Rorb* were higher in CoA, and even BLA types. LA marker *Negr1* was low in a subset of BLA, compared to LA types; but was expressed in virtually all other amygdala neurons, too. CeA GABAergic cells often followed known principles of striatal MSNs (*Ppp1r1b+, Drd1* and/or *Adora2a*), and we found refined markers that may help clarify their roles in valence encoding at greater resolution than the *Prkcd* vs. *Crh* convention. We further propose *Zfhx3* as a marker that distinguishes CeA cells from the highly related CP. Intercalated cells (ITCs) of the IA were most related to CeA cells (*Meis2+Ppp1r1b+Penk+)*. Unlike projecting inhibitory clusters of the CeA and MeA, ITCs also share expression with local inhibitory neurons (*Nfib+/Tshz2*-), providing clues on how projection distance may be encoded. One novel medial CeA cell type (GABA 21) expressed vitamin D receptor *Vdr*, along with a panel of neuropeptides. Future studies may investigate the functional significance of this unique cell type.

In the medial amygdala, one group of prolactin receptor-expressing inhibitory neurons (*Prlr)* is of particular interest for parental behavior, and several cell types we found were coherent with a recent study of *Esr1+* MeA cells (e.g., those marked by *St18, Calcr, Esr2*). The CRF-family peptide urocortin 3 (*Ucn3)* was a rare example of a single-marker gene used in functional MeA studies^45^ that indeed conformed to one discrete cell type (GABA 24). The MeA also contained many neuropeptide-enriched glutamatergic cell types of the VGLUT2 class that, to the best of our knowledge, have not been functionally addressed before. These glutamatergic types were often shared or bore resemblance to cell types found also in the basomedial amygdala (BMA). Beyond the presence of neuropeptide *Adcyap1*^65^, the BMA has not been molecularly described in the literature. We found the BMA contained a surprisingly large number of cell types. Except for the MeA, we found some overlap of these cell types with anterior and/or cortical amygdala. It would be of great interest to unravel their role in functional circuits and to determine whether they reflect the dual participation of BMA in both fear and social behaviors. In the cortical amygdala, we found distinct anterior and posterior types of *Trh* neurons, previously studied as one population, and other, novel anterior types defined by *Tgfb2*, and others. In the nucleus of the lateral olfactory tract, we identified and validated a large population of molecularly distinct *Sim1-C1ql1* neurons (GABA 8) that may be involved in associating context-based odor information with behavior^66,67^.

More generally, our data strongly support the notion that the amygdala is a somewhat arbitrary collection of regions. Amygdala neurons were highly diverse, which denoted primarily their association with a developmental subregion. At the same time, our study revealed principles underlying the organization of the regions, as well as their developmental or functional relation to one another. For instance, we described the characteristic division of glutamatergic cells into their vesicular glutamate transporters *Slc17a7* or *Slc17a6*, and other gene modules, consistent with functional divisions. Here, the cortical amygdala stood out, with distinct molecular markers expressed along its anterior-posterior axis, including class-defining vesicular glutamate transporters and branch point marker genes such as *Zeb2. Zeb2* and others (e.g., *Ptk2b, Satb1, Mafb*) also reflected organizing principles of GABAergic neurons, such as discriminating local vs. long-range projecting types.

We present the first taxonomy-wide quantification of transcriptional response to fear-learning. We argue that single-cell resolution, and unbiased, transcriptome-wide coverage is critical in identifying the response that is expected in only a small subset of cells (the engram). To this end, we applied two independent approaches utilizing the high-dimensionality of scRNA-seq. First, we compared activated, vs. non-activated cell states, per cell type. Essentially, this mimicked the experimental use of mouse models using fluorescent reporters to identify IEG-expressing engram neurons^68,69^ (e.g. *Arc::dVenus*) *in silico*. We found, however, that a single IEG was unlikely to mark the full pallet of neuronal activation at every timepoint and we opted to use a consensus panel of activity regulated genes, or IEGs instead. Indeed, different IEGs were highly expressed in just a subset of neurons and displayed time-dependent dynamics in response to fear conditioning. This observation was in line with recent reports using transcriptomics Methods^22^, where the detection of activity regulated genes expanded well beyond the brief expression windows described in the past. In our second approach we present gene expression correlation as a new, powerful tool to identify subsets of cells within cell types that display orchestrated responses related to learning and memory, especially when IEG induction is absent (recall). Independently, both approaches revealed that not all neuronal types were equally likely to participate in the response to fear conditioning. In fact, closely related molecular cell types from the same anatomical compartment could have a strong response, or completely lack activation. This observation may help explain the encoding of opposing valences (e.g. appetitive vs. aversive) in cells and circuitries of the same compartments^2,4,53^. Importantly, we also identified signatures that were regulated in activated subsets within each cell type. We confirmed known actors, and described several new candidate genes, with some class-specificity. For example, *Bdnf* and *Nlgn1* could serve as predictors for induction of learning-related cellular processes in similar datasets. Activated engram cells upregulated and coregulated activity-response genes, and processes of synaptic signaling, plasticity, development and neurite outgrowth.

Single-cell transcriptomics was used for detecting more general transcriptional signatures of neuronal activation^18,70^. It has recently been suggested and gained momentum as a tool for comprehensive and unbiased identification of cellular and molecular substrates of learning and memory^22,23,71^. As a word of caution, however, the approach and our study contain a few limitations. First, even highly refined current protocols dependent on physical dissociation of live cells may continue to suffer from dissociation stress and sampling noise, obscuring subtle expression signatures. Second, to identify subtle transcriptional states at cell-type resolution requires deep sampling, making scRNA-seq an expensive approach, especially in several experimental groups and biological repeats. As a result, we were not able to include all relevant behavioral controls, which for example does not allow us to distinguish the effect of context or cue. Finally, given the volume of data, we provide validation of neuronal types, but it was beyond the scope of the current study to validate activated states functionally.

In sum, we provide a resource of amygdala neuron populations, resolved in molecular and spatial detail. The wide scope and unbiased nature of this study revealed known and novel amygdala cell types, and their activated states. We describe cell type marker genes and gene modules regulated as a response to CFC. We provide a browsable companion website of cluster-wise scRNA-seq gene expression and spatial correlation data, for independent data exploration. Given the great knowledge about amygdala functions and circuitries, and a plethora of available behavioral paradigms and genetic tools, this taxonomy may serve to clarify the particular contributions of molecularly defined neurons, and dissect the participation of specific candidate genes in learning and memory.

## Methods

### Mice and ethics

All mice and their corresponding experimental groups are listed Table S1. We used adult C57Bl/6J wildtype mice, 7-12 weeks old; 22 males, 1 female. Most mice were of the C57Bl/6JOlaHsd subline (Envigo), and therefore carried a known alpha-synuclein (*Snca*) deletion previously evaluated with respect to fear conditioning^72^. All mice were housed under standard conditions and provided chow and water *ad libitum*. Mice were housed under a reversed day-night cycle, and all experiments were carried out during the night cycle. All experimental procedures followed the legislation under the Israel Ministry of Health - Animal Experiments Council and were approved by the institutional Animal Experiments Ethics Committees at the Technion Israel Institute of Technology and Haifa University.

### Tone-cued fear conditioning

7-11 week old male C57Bl/6JOlaHsd mice were habituated to a behavioral arena for 20min, followed by 6 cycles of US-CS pairing (cued fear conditioning, CFC): 35sec tone (CS), 750ms 0.7mV foot shock (US), 24sec rest. US-CS association was quickly and robustly established, characterized by freezing behavior during tone cue, typically after the second cycle (data not shown). For learning and consolidation, mice were sampled 2h (n=5), 8h (n=4) and 24h (n=4) after CFC. For recall, 24h after CFC, 4 mice were exposed to 6 cycles of the tone cue alone (35sec CS, 25sec rest). The 3 mice exhibiting strongest freezing behavior compared to non-FC controls undergoing the same session (video quantified, data not shown) were sacrificed 2h later. Naïve, home cage (HC) littermates served as controls in each experimental batch (n=7).

The data used for analysis of fear conditioning was collected in two experimental batches (Suppl. Table 1): Batch A was collected to examine consolidation (8h (n=4), and 24h (n=4) post-CFC, versus naïve (n=2)), while in batch B we investigated earlier responses, either 2h after CFC (n=2), or 2h after CS-recall (where CS-recall was 24h after CFC, n=3), vs. naïve (n=2). The remaining samples were used for the cell type taxonomy only.

### Perfusion

Mice were sacrificed by an overdose of Ketamine/Medetomidine, followed by transcardial perfusion with freshly prepared, ice-cold, carboxygenated NMDG-based artificial cerebrospinal fluid (aCSF: 93mM NMDG, 2.5mM KCl, 1.2mM NaH_2_PO_4_, 30mM NaHCO_3_, 20mM HEPES, 25mM D-glucose, 5mM Na-ascorbate, 2mM thiourea, 3mM Na-pyruvate, 10mM MgSO_4_, 0.5 mM CaCl_2_, adjusted to pH7.3-7.4 with concentrated HCl)^24^. Brains were quickly removed and maintained on ice in aCSF.

### Amygdala dissection

Freshly aCSF-perfused brains were quickly mounted on a Leica VT1200S Automated Vibrating Microtome and sectioned to 300μm coronal slices. Sections were then quickly microdissected in cold aCSF, to sample the full amygdala, according to the following anatomical enclosures: for medio-lateral reference, we used hypothalamus and optic tract medially; and laterally, a notional ventral extension of the amygdalar capsule of the corpus callosum fiber tracts. The beginning and end of the bifurcation of the external and amygdalar capsules marked visible anterior-posterior amygdala sampling borders. Microdissections were performed but as anatomically precise and reproducible as possible.

### Cell suspensions and scRNA-seq

Microdissected tissue pieces were digested in 800-1000μl papain digest solution per amygdala (Worthington Papain system, vial 2 reconstituted in 5ml aCSF, and 5% DNase (vial 3 reconstituted in 500μl aCSF)), 25-30min at 37°C, until mechanical trituration with a wide-diameter fire-polished glass pipette easily separated most of the tissue. Next, remaining tougher vascular or ventricular pieces were removed by filtering the digested suspension through an aCSF-equilibrated 30μm cell strainer (Partec CellTrix), to a BSA-coated microcentrifuge tube. Cells were pelleted 200g 5min at 4°C and resuspended in 200μl aCSF with 2.5% DNaseI (Worthington Papain system, vial 3 reconstituted in 500μl aCSF). For myelin and debris removal, the suspension was layered on top of 1ml 5% OptiPrep (Sigma) in aCSF in a BSA-coated microcentrifuge tube, and centrifuged for 6min 150g at 4°C, with slow ramping. The resulting cell pellet was resuspended in a minimal volume of aCSF, and inspected in a Burker counting chamber for intact cell morphologies and high viability and successful debris removal. At all steps, from perfusion to final single cell suspension, tissue or cells were maintained in ice-cold carboxygenated (95% O2, 5% CO2) aCSF – with the exception of papain digest, where the temperature was 37°C.

Single-cell suspensions were diluted to 1,000 cells/μl, and processed for 10×Chromium-v3 GEM generation and scRNA-seq. We followed the manufacturer’s instructions, targeting 5,000-6,000 cells per sample. Sequencing libraries were multiplexed and sequenced on Illumina NextSeq or NovaSeq NGS platforms, targeting a depth of >35-40K reads per cell.

### Quantification and statistical analysis of scRNA-seq data

To interpret our scRNA-seq data for the purpose of cell-type discovery, we largely followed the logic of our tested-and-proven analysis pipeline, detailed in^17^. First, raw sequencing data were demultiplexed, aligned with the genome, and mRNA molecules counted on the *cellranger* pipeline (10x Genomics), resulting in output matrix UMI count files. Next, we conducted informal exploratory analysis and found robust division of cells into non-neuronal and three main neuronal classes (GABA, VGLUT1, VGLUT2), which led us to apply the following iterative clustering approach.

#### Iterative clustering

All final analysis was performed in our MATLAB-based clustering pipeline (described in the next section), in the following three iterations:

##### Step 1: Cell QC and classification

To extract and classify neurons, we set the threshold of 3,000 UMI/cell and 2,500 genes/cell, then follow standard pipeline steps (see below) to obtain rough clusters that were classified into four categories (GABA, VGLUT1, VGLUT2, non-neuronal) based on a majority vote of known markers (*Gad2*=GABA, *Slc17a7*=VGLUT1, *Slc17a6*=VGLUT2 and non-neuronal cells (which were excluded) = *C1qc+C1qa+C1qb+Gja1+Cx3cr1+Acta2+Ly6c1+Mfge8+Plxnb3+Cldn11+Aqp4+Vtn+Cldn5+Pdgfrb+Flt 1+Slc25a18+Pdgfra+Foxj1+Olig1+Olig2+Sox10+Hbb-bs+Hbb-bt+Hba-a2+Ttr*). Doublets were called and excluded at this point if a cluster combined any of the above gene sets at a ratio <2, except the permitted combination of VGLUT1 and VGLUT2.

To identify and classify non-neuronal cells, we set a lower threshold of 2,000 UMI/cell and 1,000 genes/cell, and excluded cells that expressed either *Gad2*, *Slc17a7*, or *Slc17a6*. Non-neuronal clusters were classified into the following categories: immune (based on *C1qc, C1qa, C1qb, Mrc1, Pf4, Cx3cr1*), astrocytes (based on *Gja1, Aqp4, Foxj1, Aldoc, Mfge8, Slc25a18*), vascular (based on *Vtn, Cldn5, Pdgfrb, Flt1, Acta2, Ly6c1*), oligodendrocytes (based on *Plxnb3, Cldn11, Olig1, Olig2, Sox10, Pdgfra*), and blood (which were excluded, based on *Hbb-bs, Hbb-bt, Hba-a2, Ttr*).

##### Step 2: Cluster calling, QC, and doublets removal

For retained cells of each Step 1 category (GABA, VGLUT1, VGLUT2, Immune, Astrocytes, Vascular, Oligodendrocytes) we reran separately the standard pipeline, which produced more class-relevant gene sets during feature selection. The resulting putative cluster lists and visualizations aided manual inspection to (a) exclude remaining suspected doublet-clusters and (b) merge clusters with extremely high similarity to neighboring clusters. This step resulted in the final cluster list. We then assigned a two-gene identifier name to each neuronal cluster, based on highly enriched genes and a literature survey.

##### Step 3: Final feature selection and visualization

After Step 2 doublet removal and merging of similar clusters, we repeated only feature selection and replotted all visualizations, as presented in the study.

#### Standard clustering pipeline scRNA-seq

In each of the steps outlined above, we applied the following:

##### Normalization

Each cell (vector) was normalized to a length of one, then multiplied by 20,000.

##### Gene exclusion

We excluded 53 immediate early genes (IEGs: *Btg2, Jun, Egr4, Fosb, Junb, Gadd45g, Fos, Arc, Nr4a1, Npas4, Coq10b, Tns1, Per2, Ptgs2, Rnd3, Tnfaip6, Srxn1, Tiparp, Ccnl1, Mcl1, Dnajb5, Nr4a3, Fosl2, Nptx2, Rasl11a, Mest, Sertad1, Egr2, Midn, Gadd45b, Dusp6, Irs2, Plat, Ier2, Rrad, Tpbg, Csrnp1, Peli1, Per1, Kdm6b, Inhba, Plk2, Ifrd1, Baz1a, Trib1, Pim3, Lrrk2, Dusp1, Cdkn1a, Pim1, Sik1, Frat2, Dusp*5), sex genes (*Xist, Tsix, Eif2s3y, Ddx3y, Uty, Kdm5d*), and genes that were not relevant (for example, when clustering neurons, all non-neuronal markers mentioned above were excluded).

##### Feature selection

First, we selected only genes with expression in at least 5 cells, but less than 50% of all cells. We then used CV vs. mean fit to rank genes as described before^73^. The number of genes selected was decided based on plotting the distance to the fitted line, from the largest to smallest distance, and finding the “bending” point closest to the origin.

##### PCA

Next, we calculated PCA projections, deciding the number of PCs based on the optimal point of explained variance vs. PC.

##### Batch correction

Because data collection was performed in three time-separated batches (Suppl. Table 1), we corrected for batch effects applying the HARMONY algorithm^74^ to the PCA projection.

##### Dimensionality reduction by t-SNE

Two-dimensional embedding was performed using t-SNE (MATLAB implementation) with correlation distance, Barnes-Hut algorithm, *theta*=0.5, learning rate=#cells/12, exaggeration=20. The number of PCs was as determined above. To choose perplexity, we used the following heuristics: on PCA projections, we calculated the distance (correlation) of each cell to its first 500 neighbors and determined the optimal “cutting” point. This produced an “optimal” number of neighbors per cell, and we chose the perplexity to be the median of this vector. The t-SNE was initiated based on the first two dimensions of the PCA, as described in^75^. For t-SNE visualization of the final clusters (Step 3, as shown in the figures of the article), we used Perplexity=100 and Exaggeration=5.

##### Clustering

Next, we used the DBSCAN algorithm to cluster the t-SNE-embedded cells. We chose DBSCAN parameters based on visual inspection of the resulting clusters, aiming to avoid over-clustering, without losing the sensitivity to detect small clusters.

##### Post-clustering

After clustering, we removed cells that DBSCAN labeled as outliers, and sorted the cells in each cluster using 1D t-SNE (applied to each cluster separately).

##### Dendrogram construction

To build the cluster dendrograms (a 1D order of clusters), first, we log2(x+1)-transformed the expression; second, calculated cluster-wise mean expression profiles; third, calculated the PCA projection of the matrix; fourth, used the projection coordinates for linkage clustering (Ward algorithm, correlation distance); and finally, used the MATLAB function “optimalleaforder” to order the clusters.

##### Branch point marker gene identification

To help us

a. identify genes enriched in multiple clusters and
b. define main splits along the cell type hierarchy, we computed branch point marker genes. At each split along each class dendrogram, we calculated the difference of fraction positive cells on either side of the branch. The resulting top scoring genes, on either side, were considered branch point markers.

### Allen Mouse Brain ISH Atlas-based spatial correlation analysis

To align the gene expression profiles detected for each scRNA-seq-derived cluster with its spatial context, we performed correlation to the Allen Mouse Brain *in situ* hybridization Atlas (mouse.brain-map.org)^25^, as we described in^17^. Briefly, we used the AMBA aggregated data, which provides an “energy score” for each gene from the *in situ* experiment, per each voxel of 200×200×200μm. Each 200μm voxel also has an AMBA-generated brain region annotation (the Atlas has a finer resolution of 25μm voxels). Next, we calculated the *rho* correlation of each cell type (average expression profile) and each voxel in the brain (left hemisphere), as follows: we defined genes that have valid expression in the AMBA database as covering >30 voxels, a >5 “energy score”, and an average over all voxels >0.2. The set of quality genes was intersected with the set of genes from the final feature selection of our scRNA-seq data analysis (Step 3). The *rho* correlation of each cell type with the region-annotated 200μm voxels is presented in Supplementary Table 3. For better resolution and smooth visualization, we parcellated the coarse 200μm voxels to a finer, linear interpolated 25μm grid, and visualized resulting heatmaps on one coronal hemisphere.

### AMBA-based virtual in situ hybridization (Slc17a7, Slc17a6, Gad2)

Based on the AMBA histology *in situ* hybridization (ISH) gene expression database, we sampled three representative coronal sections, spanning the anterior-posterior amygdala for three neuronal marker genes (9 sections overall): *Gad2* for inhibitory neurons, and *Slc17a7* (VGLUT1) and *Slc17a6* (VGLUT2) for excitatory neurons. We segmented ISH gene expression to cells using global thresholding. We aligned the sections with the AMBA for region annotation, accounting for section distortion along the dorsoventral and mediolateral axes. Next, we normalized expression levels for each gene in all aligned section subregions, to uncover differential levels of expression relative to subregion size and visualized them as heatmaps.

### Spatial transcriptomics (ST) Visium (10x Genomics)

For spatial transcriptomics, 2 adult mice (1 male, 1 female) were sacrificed by transcardial perfusion with aCSF. We quickly extracted, coated, and cryomold-embedded the fresh brains in cryoprotective OCT (TissueTek), flash-froze them in isopentane equilibrated on dry ice, and maintained them in sealed bags at −80°C until processing. Per brain, we collected 4 right-hemisphere coronal cryosections at 10μm thickness, aiming at approximately 200-300μm spacing, spanning the anterior-posterior axis of the amygdala, onto the Visium ST gene expression slide (4 capture areas, 10x Genomics). For tissue preparation, we followed the manufacturer’s instructions, with the following specifications: methanol fixation, immunofluorescence staining with DAPI only (no antibody), imaging at 4× on a Nikon Eclipse Ti2: DIC, DAPI, and TRITC channels for fiducial and section alignment and 25min permeabilization, as we had previously determined on amygdala test sections, using the Tissue Optimization kit. We then proceeded with the Visium Gene Expression Kit following the manufacturer’s instructions, with 15 PCR cycles for cDNA amplification. Sequencing was performed on Illumina NGS platforms to a depth of 150-200M reads per sample (i.e., capture area, or section).

### Visium ST alignment and quality control

To map the anatomical annotation for each 2D capture spot, we aligned the DAPI images of Visium ST sections to the AMB Reference Atlas, correcting for distortion caused by sectioning along the dorsoventral and mediolateral axes. Sections were on average 100μm apart, with considerable variability (z-axis resolution); mRNA-capture spot diameters were 55μm, centers 100μm apart (x-y resolution). An average 3,226 capture spots covered each coronal hemisphere, 5-10% of which mapped to the amygdala (mean 272 spots). Spots that aligned with the amygdala contained ~2,500-3,500 genes and ~30,000 UMIs.

### Quantification and statistical analysis of Visium ST data

With the exception of the spatial barcodes replacing cell barcodes (spots instead of cells), analysis was similar to the scRNA-seq analysis pipeline described above. Each spatial barcoded spot of 55μm diameter was expected to capture multiple cells (including neurons, glia, and vasculature), resulting in “micro-bulk” expression data. We previously found non-neuronal cells to be less spatially distinct in the brain^17^, and therefore, expected genes derived from non-neurons to be noisy, sporadic, and spatially less informative. To focus on neuronal diversity instead, we removed non-neuronal marker genes from analysis.

Briefly, after preprocessing, filtering, normalizing, and feature selection, we performed dimensionality reduction by principal component analysis (PCA) of the high-dimensional expression data, followed by 2D embedding with t-distributed stochastic neighbor embedding (t-SNE), based on their similarity in the high-dimensional gene expression space. This resulted in a 2D map. Focusing on neuronal markers, we clustered these spots according to their distance in 2D, using the density-based spatial clustering of applications with noise (DBSCAN) algorithm. Thus, spots with similar gene expression were grouped together in the same cluster. We also used *k*-nearest neighbors (KNN) to regroup the outlier spots obtained from the DBSCAN algorithm into their closest neighbor cluster. Next, we remapped the spots to their original location on the tissue, based on the spatial barcode index, and inspected whether the clusters followed an unbiased spatial distribution in the tissue.

### Visium ST-based spatial correlation analysis

Analogously to AMBA-based spatial correlation (above), we used Visium ST expression data to infer the spatial distributions of scRNA-seq annotated cell type mean cluster expression. First, we performed feature selection of highly variable genes for scRNA-seq and Visium ST datasets separately, then combined and intersected the two gene lists. After normalizing both datasets, we conducted pairwise linear Pearson correlations between each scRNA-seq cluster and each Visium spatial spot (2D). This resulted in a matrix of *rho* correlation values for each spot with each cell type, which we mapped back to the original xy-position of the relevant section. For 1450 global pattern comparison, we normalized the *rho* correlation values and presented all the section spots together in a heatmap form. Having a registered anatomical annotation for each 2D capture spot (see above, Spatial Transcriptomics (ST) Visium), we quantified spatial correlations for each scRNA-seq cluster per region, as presented in Suppl. Table 4.

### Multiplexed fluorescence in situ hybridization

Following the same procedure as for spatial transcriptomics (10x Visium), we extracted aCSF-perfused brains, coated and cryomold-embedded them in cryoprotective OCT (TissueTek), flash-froze in isopentane equilibrated on dry ice, and maintained in frozen brains in sealed bags at −80°C until processing. We collected 16μm coronal cryosections spanning the amygdala on 2% APTES silenized glass slides^76^, proceeded with quick post-fixation in 4% PFA for 10min, 2 washes in PBS, dehydration in isopropanol, and stored slides in 70% ethanol at 4°C until further processing. Before hybridization, sections were briefly dehydrated in 100% ethanol, airdried, and encircled with a hydrophobic barrier pen. Starting with Protease 4 pre-treatment, we used the RNAScope Fluorescent Multiplex (3-plex) Reagent Kit (ACDBio) and followed the manufacturer’s instructions. The following mouse probes were combined for 3-plexing in alternating channels: Slc17a6 #319171, Slc17a7 #416631, Gad2 #439371, Gal #400961, Avp, O2 #472261, Sim1 #526501. Sections were imaged on a Nikon Eclipse Ti2 epifluorescence microscope at 4× and 20× magnification. Image processing was carried out using NIS Elements software (Nikon), and final images, LUTs 95th to 99th percentile, were batch exported using MATLAB. To map sections to anatomical annotations, we aligned DAPI images with the AMB Reference Atlas, adjusting for distortions along the dorsoventral and mediolateral axes caused by sectioning.

### Cell activity score by high IEG expression (IEG score)

We defined a set of eight immediate early genes (IEGs): *Arc, Bdnf, Btg2, Fos, Fosl2, Homer1, Npas4, Nr4a1*, calculated the 90^th^ percentile expression for each. A cell was called activated, or IEG^high^, if it expressed any of the genes above this 90^th^ percentile. For every cell type, we calculated the fraction of activated cells per timepoint (8×5 matrix). We then summed the fraction of all eight genes per timepoint (1×5 vector) and defined the activity score as the maximum difference between CFC-samples (2h, 8h, 24h, recall) to HC control.

### Differential gene expression of activated neurons (IEG^high^)

Based on the same set of eight IEGs (*Arc, Bdnf, Btg2, Fos, Fosl2, Homer1, Npas4, Nr4a1*), we consider a cell “activated” or IEG^high^, if its expression is in the top 5% of the gene expression distribution for any of these IEGs (95^th^ percentile over all cells). For all CFC-sampled cells per cell type, we then performed differential gene expression analysis for IEG^high^ vs. IEG^low^ cells. We calculated the average expression of each group and plotted the difference vs. average (Fig. 6g). For each cell type, we registered the value of the difference (*d* = IEG^high^ – IEG^low^), and ranked genes by their cluster frequency of differential expression (*d* > 0.5) in their respective class (GABA, GLUT). Gene ontology (GO) terms enriched IEG^high^ for IEG^low^ cells per class were analyzed in DAVID, biological processes BP5 (Suppl. Table 5).

### Analysis of CFC-associated gene expression correlation

We defined seven groups of genes indicated in the literature to be related to neuronal activity-dependent transcription (Suppl. Table 6): IEG, learning/memory, Glutamate receptors, GABA receptors, K^+^-channels, Ca^2+^-channels and semaphorins. We included genes expressed in more than 200 cells per analyzed class, and cell types with ≥20 cells per sampling time point. Per cell type, per time point, we then calculated Pearson pairwise correlation for each gene list, and all gene lists combined (Correlation summary). For each cell type and time point, we next calculated correlation scores over all pairwise p-values (*score* = −log_10_ *p*), (Fig. 7d, Suppl. Table 6). The per-cluster correlation score (Fig. 7c) was the difference between the maximum score among CFC (2h, 8h, 24h or recall) – the maximum score of home cage (HC) control (batch A or B). For *Nlgn1* and *Bdnf* (GLUT only), we calculated Pearson pairwise correlations (per time point, per cluster), with all genes expressed in >5 cells, and ranked correlated genes by their frequency of correlation (p<0.01) in CFC. Gene ontology (GO) terms enriched among the top 200 correlated genes were analyzed in DAVID, biological processes BP5 (Suppl. Table 6).

## Data and code availability

All data and custom code used to perform analysis will be made available upon publication.

## Author Contributions

Study design: HH, SN, SW and AZ designed the study and planned experiments

Data collection: SN, NR, SS and MT performed fear conditioning, HH and NR performed cell preparations, OO performed scRNA-seq and Visium ST, ZL performed Visium ST, HH and MT performed in situ hybridizations.

Data analysis: AZ, HH analyzed scRNA-seq data, AZ, MT, HH analyzed spatial data (Visium ST, AMBA volumetric data, in situ hybridization).

Data interpretation: HH, AZ, SN and SW critically discussed and interpreted scRNA-seq data, HH and AZ interpreted spatial data Writing: HH wrote the paper with help from AZ and SW, and input from all authors

## Acknowledgements

A.Z. is supported by the Milgrom Family Fund, European Research Council (TYPEWIRE-852786), Human Frontiers Science Program (CDA-0039/2019-C) and Israel Science Foundation (2028912). H.H. is supported by the Swedish Brain Foundation (Hjärnfonden) and Human Frontiers Science Program (CDA-0039/2019-C). S.W. is supported by the Milgrom Family Fund. We thank Raphael Lamprecht for critical discussion of the data, and Walla Odi for technical assistance.

