## Supplemental Figures for "Cell types in the mouse amygdala and their transcriptional response to fear conditioning"

#### Extended Data Figures

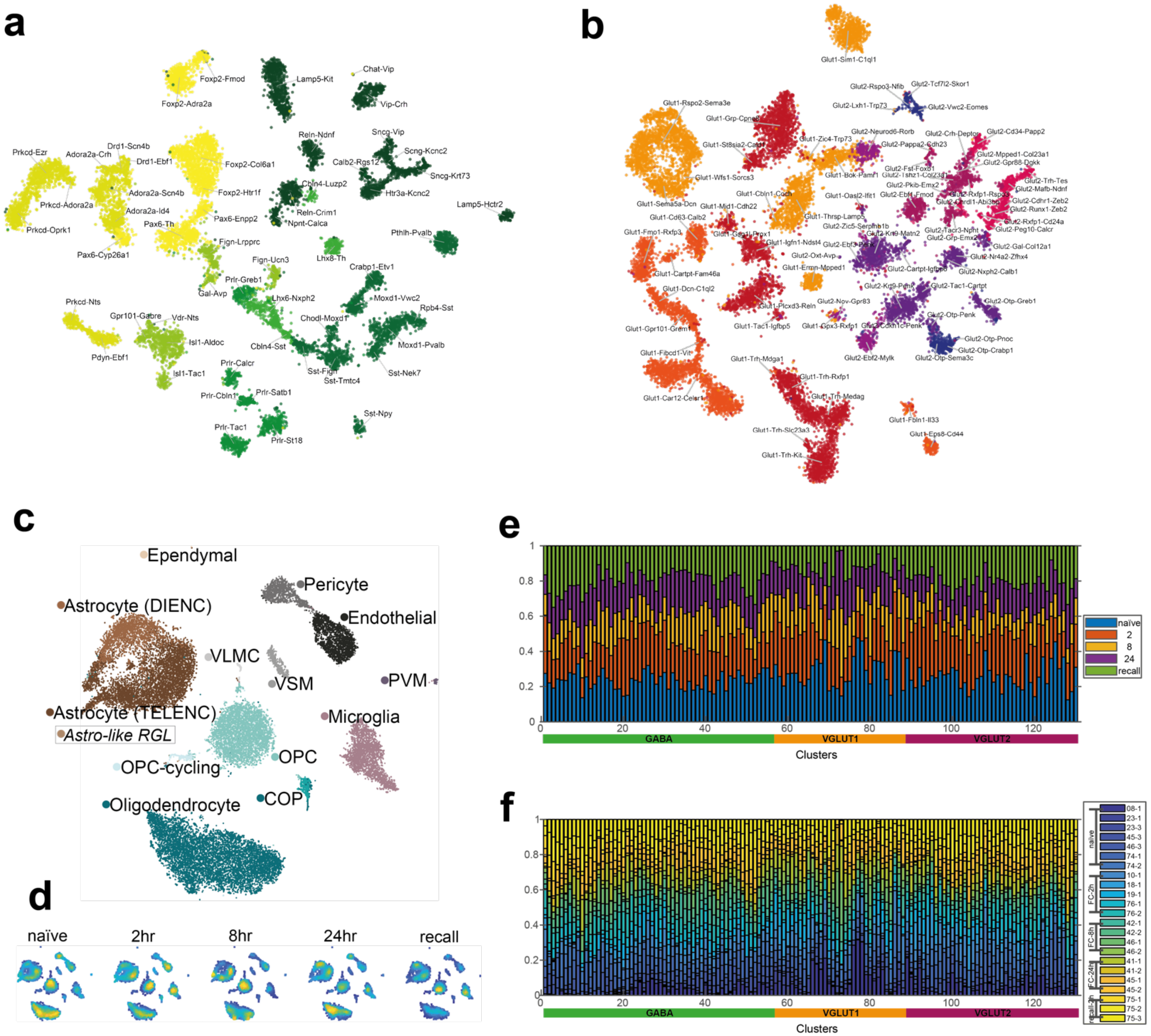


**Extended Data Figure 1** – related to Figure 1 | **Molecular cell types of the amygdala, from adult, behaving mice. a** tSNE embedding of 13,006 GABA cells, colored by clade, clusters labelled with two-gene identifier name in cluster center. **b** tSNE embedding of 17,178 glutamatergic cells (VGLUT1, VGLUT2 combined), colored by clade, clusters labelled with two-gene identifier name in cluster center. **c** tSNE embedding of 25,330 non-neuronal cells, colored by cluster (14 clusters). **d** Non-neuronal tSNE embedding as **c**, divided by experimental group. **e** Fraction of cells for 130 neuronal clusters, by experimental group. **f** Fraction of cells for 130 neuronal clusters, by sample.

### Extended Data Figures

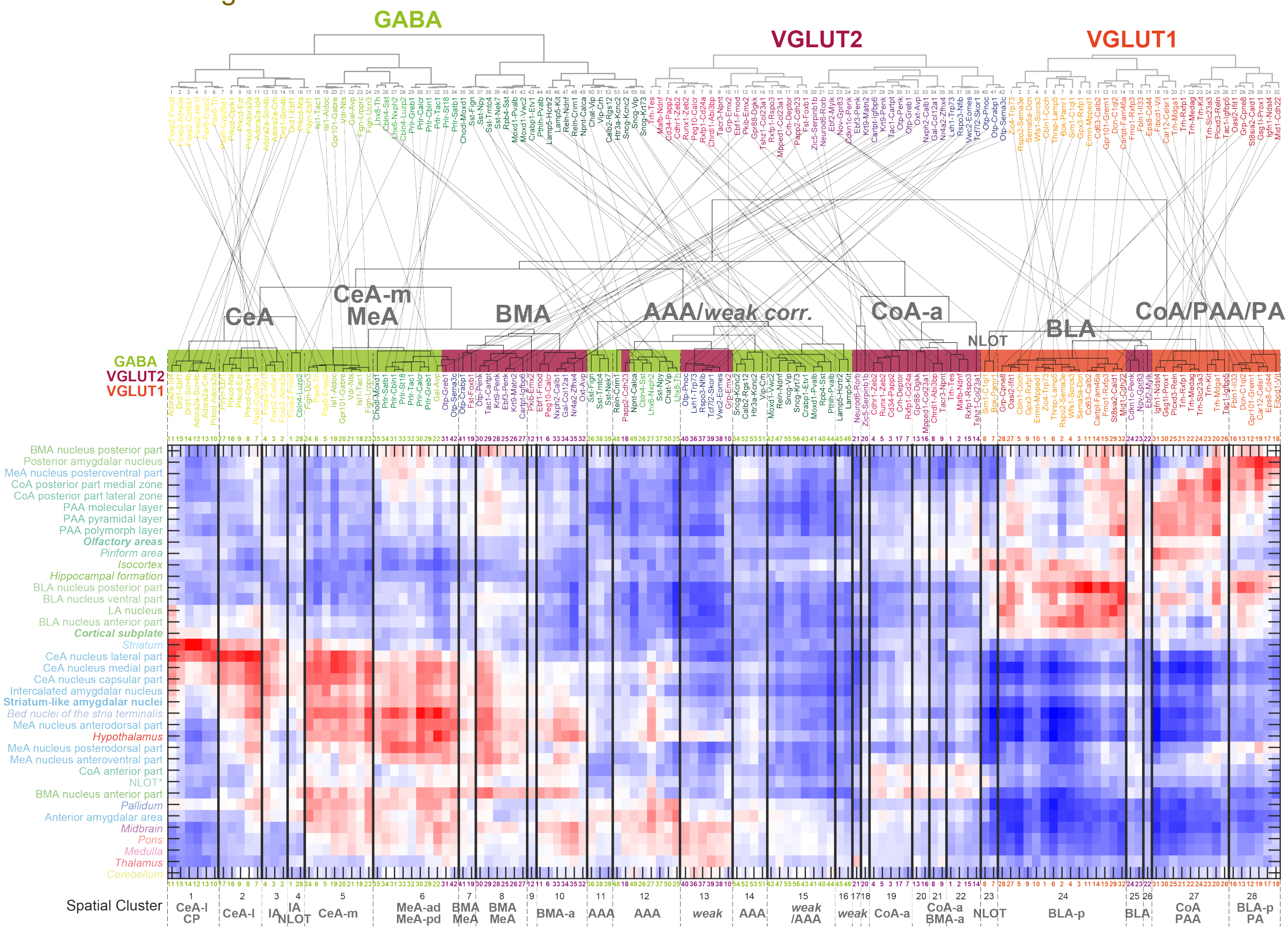


**Extended Data Figure 2** – related to Figure 2 | **Clustering of cell types by similarity of inferred spatial distributions.** Heatmap and dendrogram of cell type correlation with AMBA volumetric data (Fig. 2), columns (cell types) are arranged according to similarity in correlation with analyzed regions (rows). The dendrogram is colored by cell class, vertical lines indicate approximate "spatial clusters", i.e. cell types most strongly correlated in their spatial enrichment. Highest correlating regions indicated below. Top, for reference, scRNA-seq dendrograms of GABA, VGLUT1 and VGLUT2 classes (as Fig. 1); cell types are linked to reveal the relation of their scRNA-seq and spatial clustering.

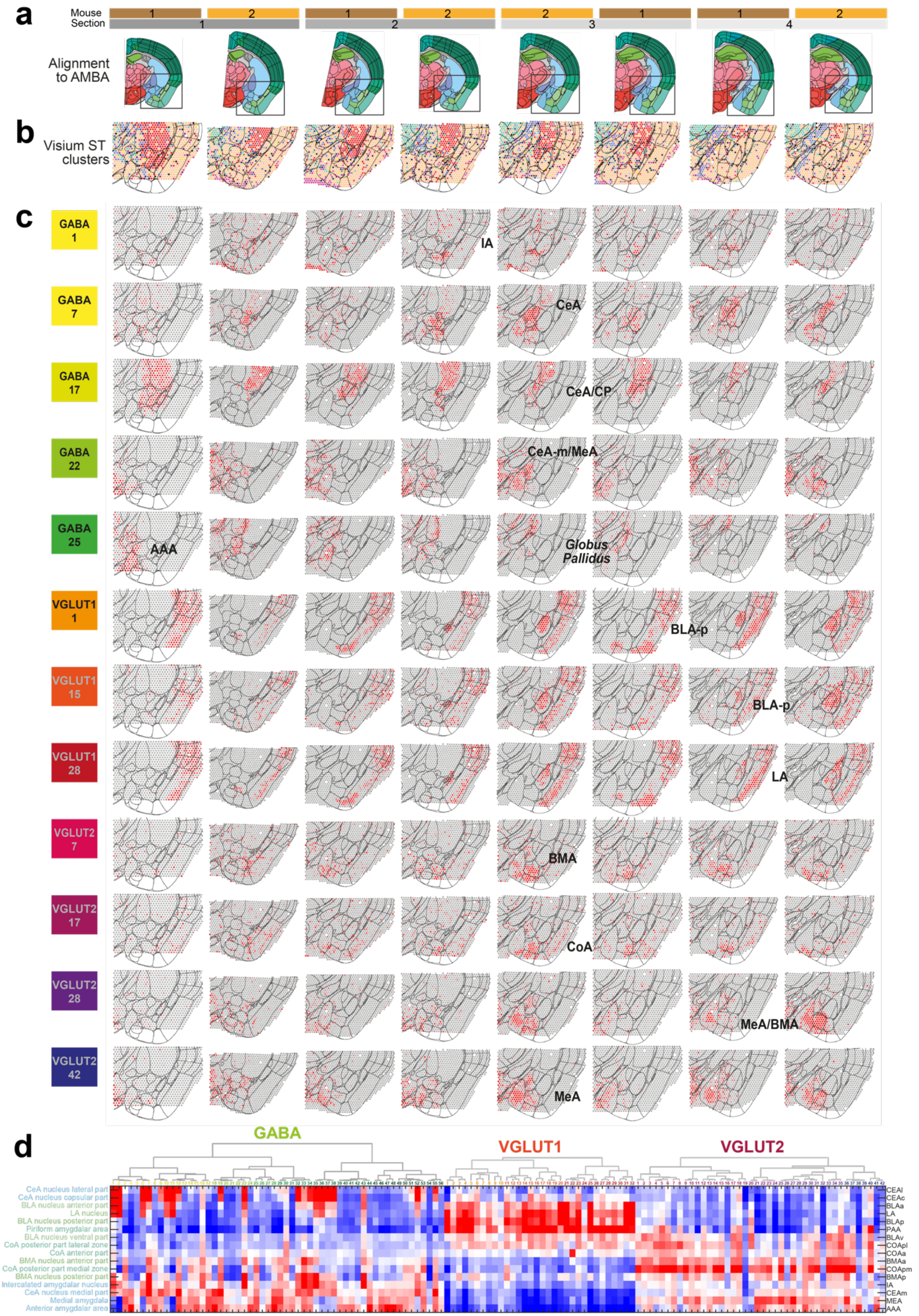


**Extended Data Figure 3** – related to Figure 2 | **Spatial transcriptomics (Visium) confirms patterns of cluster enrichment. a** Sample and section overview. 4 coronal sections covering the anterior-posterior amygdala were collected from 2 mice (1 male and 1 female), each;

*(Extended Data Figure 3, continued)* 3D alignment of all 8 sections to AMBA for anatomical reference, below. **b** Clustering of 2D Visium ST capture spots; visualized by cluster on the sections (zoom-in). Each dot represents a capture spot; colored by cluster identity. **c** Correlation heatmap between spatial transcriptomics and 12 cell types (examples), visualized on the sections, as annotated in **a**. Grey, low; red, high correlation. **d** Correlation heatmap between spatial transcriptomics expression annotated to amygdala regions (rows, ordered by similarity) and all 130 cell types (columns, as Fig. 1). Blue, low; red, high correlation. Top, dendrograms for reference.

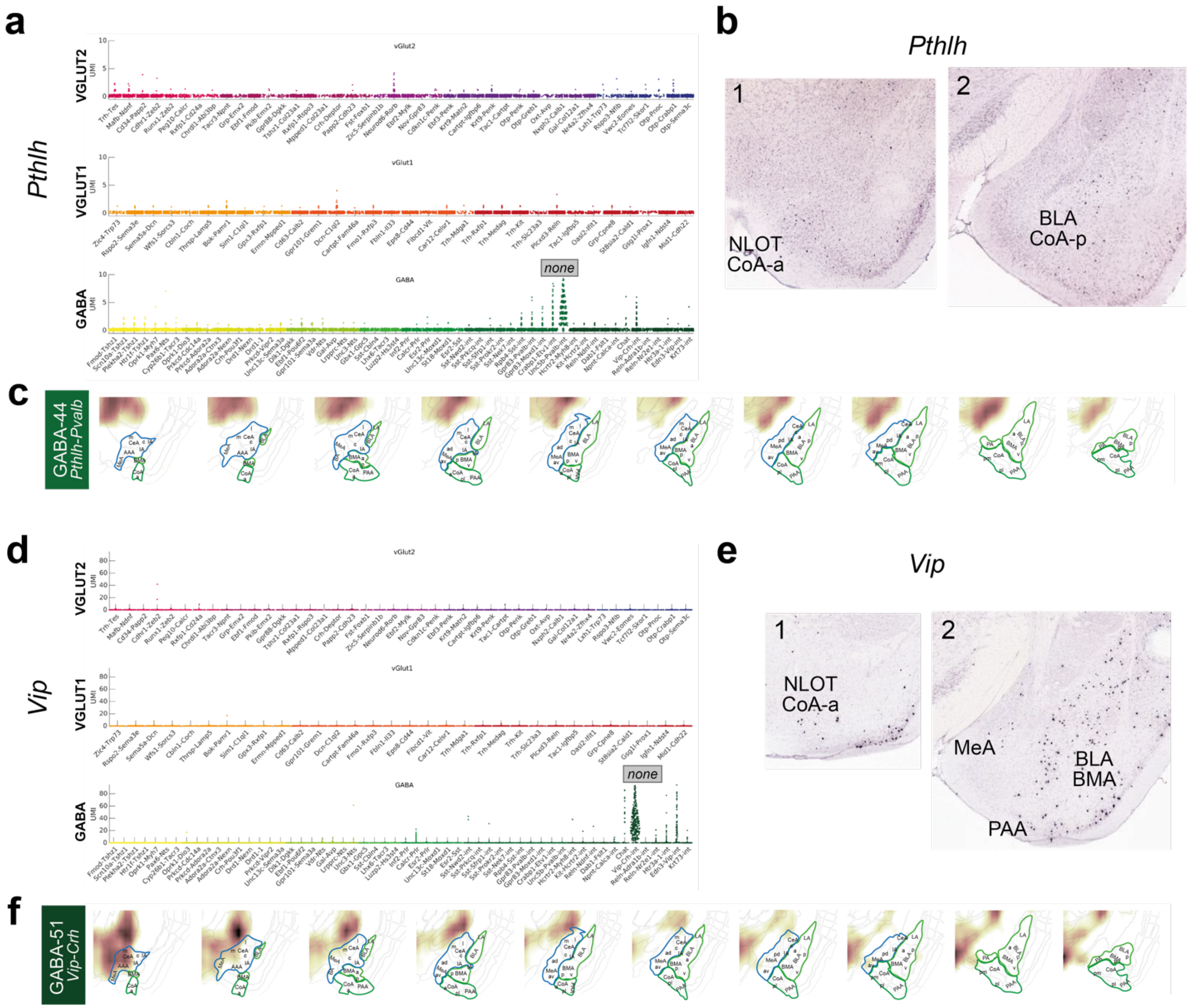


**Extended Data Figure 4** – related to Figure 3 | **GABAergic interneurons are sparse and dispersed across several amygdala nuclei. a** *Pthlh* is a marker uniquely expressed in GABAergic interneurons, especially cluster GABA-44 *Pthlh-Pvalb.* **b** AMB ISH of *Pthlh* expression in two sections of the amygdala, appears sparse and dispersed. **c** Expression correlation with *Pthlh*-cluster GABA-44 is absent in amygdala regions. **d** *Vip* is a marker uniquely expressed in GABAergic interneurons, especially cluster GABA-51 *Vip-Crh.* **e** AMB ISH of *Vip* expression in two sections of the amygdala, appears dispersed. **f** Expression correlation with *Vip*-cluster GABA-51 is weak or absent in the expected regions (e.g. BLA/BMA); correlation of the cluster to AAA is not reflected by *Vip*-ISH (**e**).

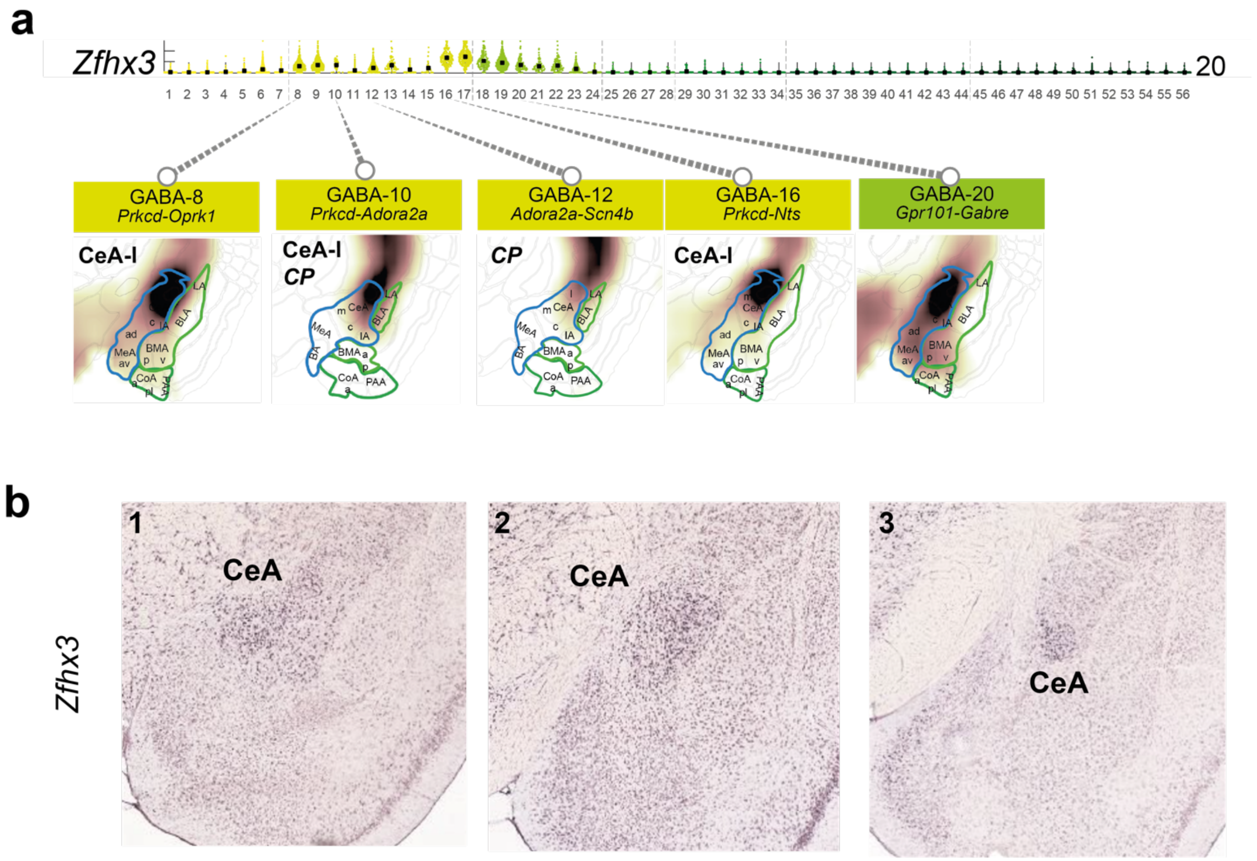


**Extended Data Figure 5** – related to Figure 3 | ***Zfhx3* expression as a marker for CeA, rather than CP, origin. a** Top: expression of *Zfhx3* in GABA neurons (y-scale maximum, 20 molecules); bottom: relevant examples of correlation with GABA clusters expressing different levels of *Zfhx3.* Correlation coefficients visualized as a heat map on coronal sections; white, low; brown, high correlation. **b** AMB ISH of *Zfhx3* showing enriched expression in the CeA, compared to surrounding and related structures, such as dorsally neighboring striatal CP.

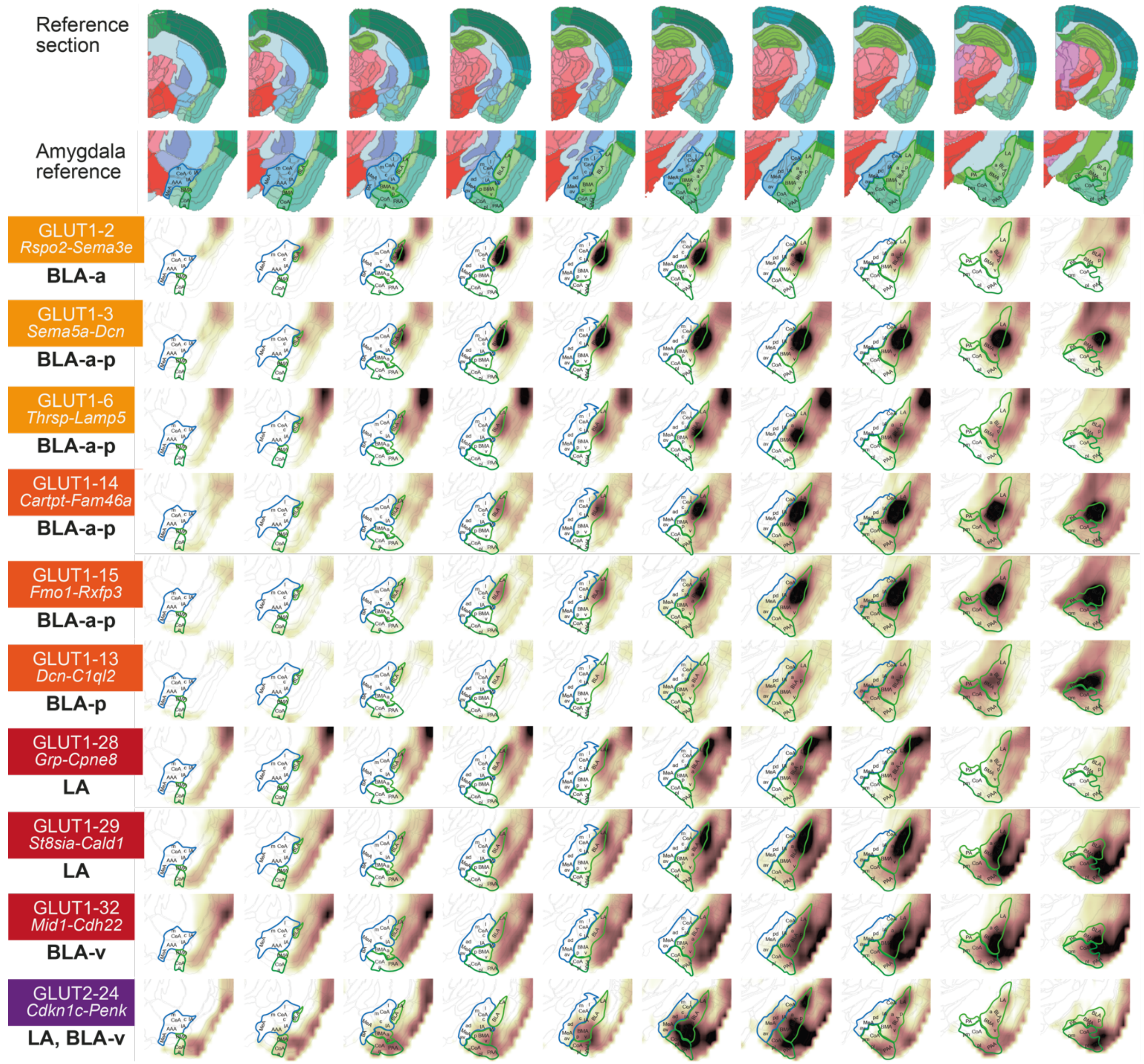


**Extended Data Figure 6** – related to Figure 4 | **Inferred spatial distributions of BLA/LA glutamatergic neuron populations using AMBA volumetric expression ISH data.** Correlation coefficients visualized as a heat map on coronal sections; white, low; brown, high correlation. Top rows, AMBA reference section, and annotated zoom-in; left column, cluster name, and most enriched structure.

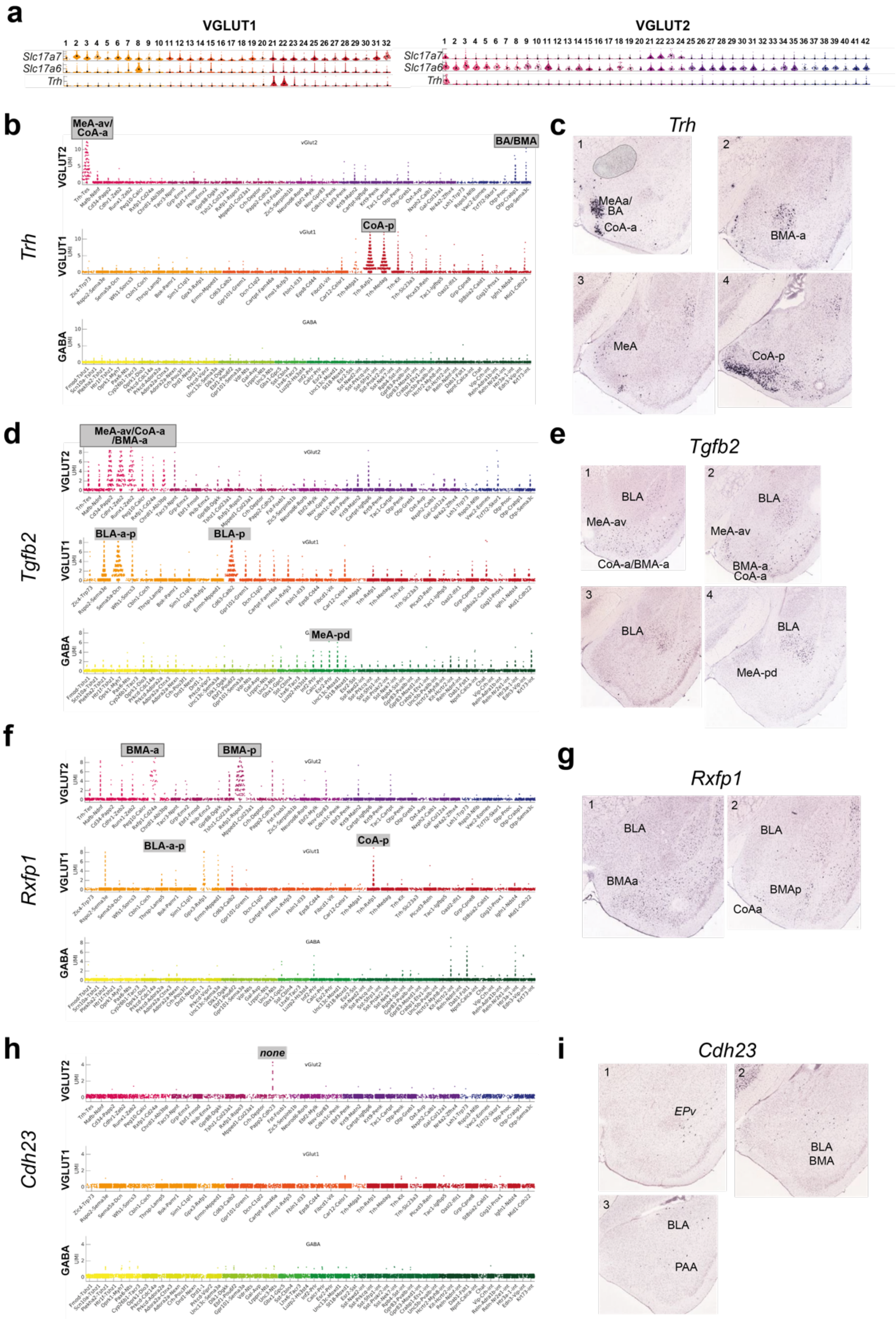


**Extended Data Figure 7** – related to Figure 5 | **Glutamatergic neurons of the amygdala's olfactory-related regions. a** *Trh+* glutamatergic populations coexpress VGLUT1 (*Slc17a7)* and VGLUT2 (*Slc17a6*). **b** *Trh* expression is restricted to few glutamatergic types. **c** AMBA ISH of *Trh* on 4 sections, with amygdala regions expressing *Trh* highlighted. **d-e, f-g, h-I,** all as for **b-c**, but for marker genes *Tgfb2, Rxfp1* and *Cdh23*; some of which mark clusters that were not validated using full spatial correlation analysis.

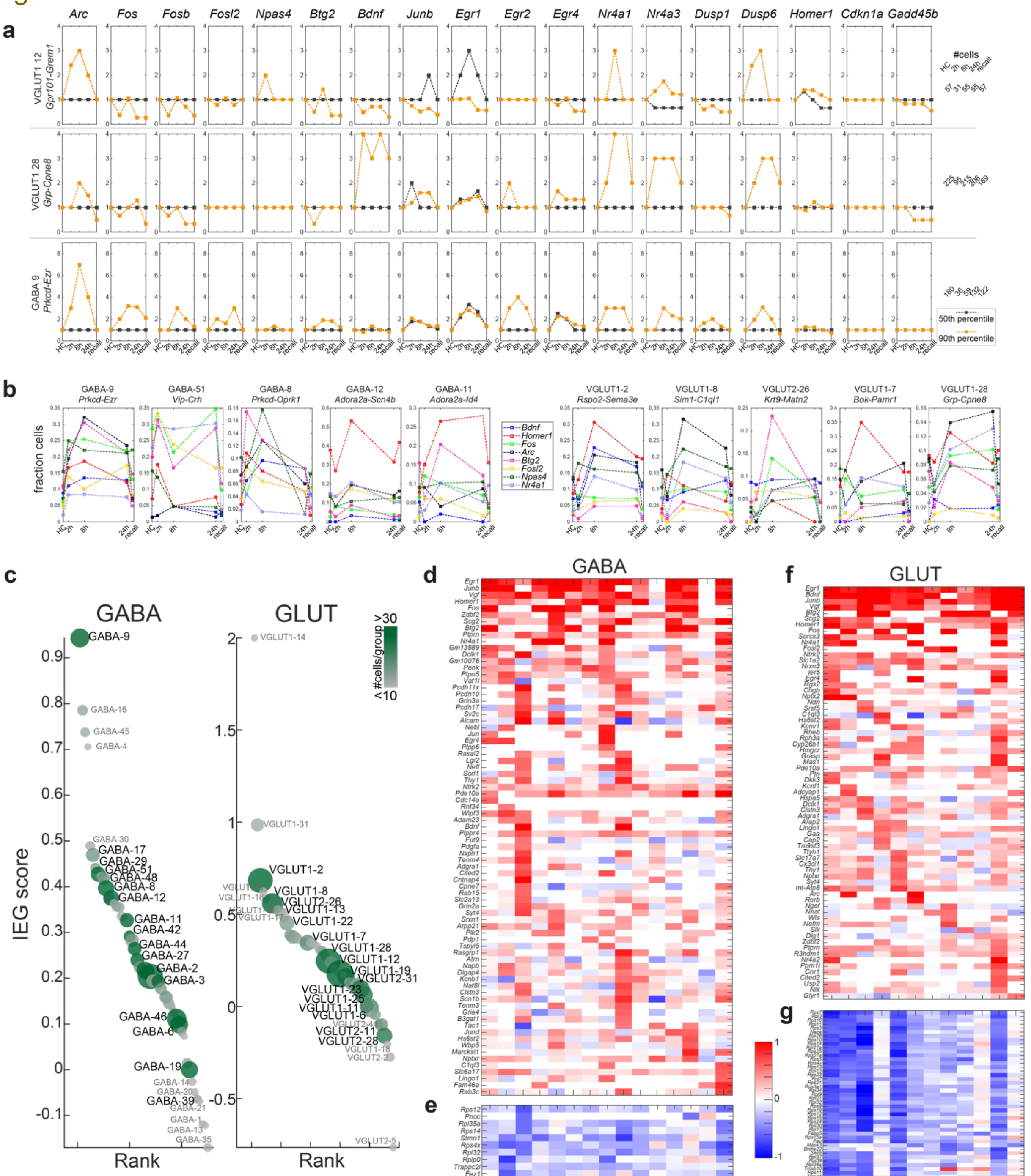


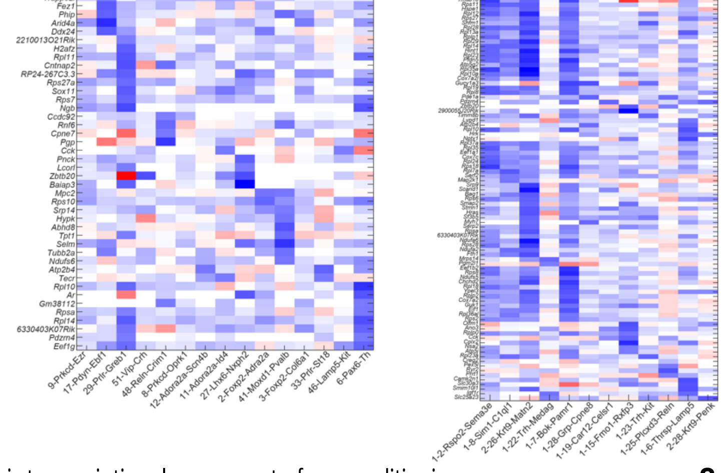


**Extended Data Figure 8** – related to Figure 6 | **Activated cell states across the neuronal taxonomy**. **a** Time-resolved expression of IEGs for three cell types; median, black; 90th percentile, orange. **b** As Fig. 6d, in fraction cells expressing in the 90th percentile, eight IEGs used for IEG score, in the five top IEG-scoring GABA and GLUT types. **c** GABA and GLUT clusters ranked according to IEG score. Circle size represents cluster size, color represents size of smallest sampling group (grey; <10 cells, dark green; >30 cells). **d-g** Heatmaps of upregulated (**d, f**) and downregulated (**e, g**) genes in IEG+ cells, per cluster, large clusters only. Cluster order by IEG score. Blue, low; red, high expression.

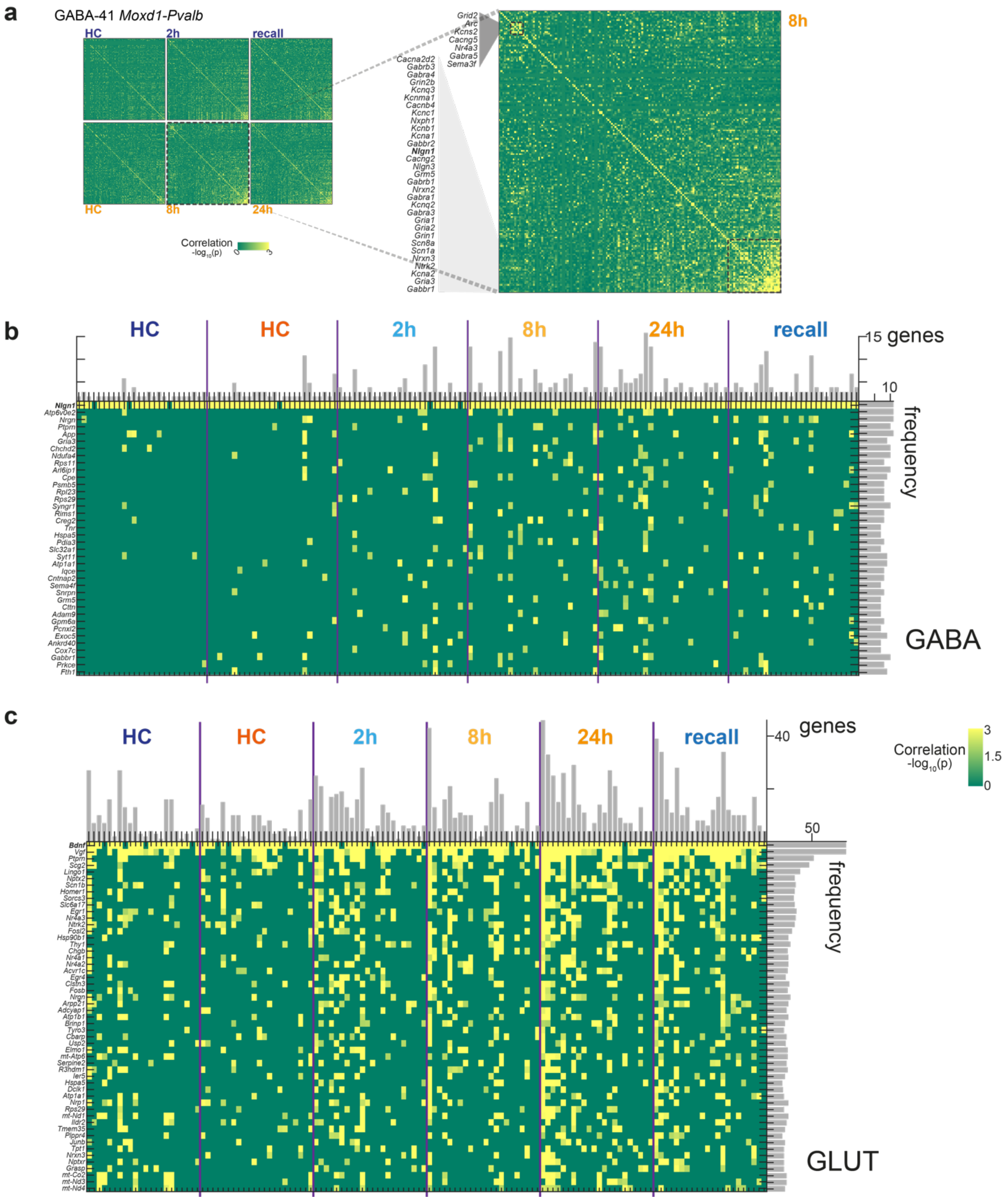


**Extended Data Figure 9** – related to Figure 7 | **a** For GABAergic interneuron type *Moxd1-Pvalb,* pairwise correlation of 155 learning-related genes (rows and columns), per post-CFC sampling time point and batch-specific home cage (HC) control. Pearson coefficient (green, low; yellow, high), genes are ordered by hierarchical clustering. Zoom-in to 8h post-CFC (right), with two correlated gene expression modules highlighted. **b-c** For large clusters, Pearson coefficient for genes highly and frequently correlated with neuroligin *Nlgn1* in GABA (**b**) or *Bdnf* in GLUT (**c**), per sample group (timepoints post-CFC, and naïve home cage (HC) per batch (orange, batch A; blue, batch B). Green, low; yellow, high. Rows, genes; columns, clusters (ordered by overall correlation score, as Fig. 7e). Bars on the top and left are the sum of each column or row (p>0.01), respectively.
